# FcγR- and CD9-dependent synapse engulfing microglia in the thalamus drives cognitive impairment following cortical brain injury

**DOI:** 10.1101/2024.09.19.609743

**Authors:** Ken Matoba, Takahiro Kochi, Yassin R Mreyoud, Jana H. Badrani, Hency Patel, Hiroshi Tsujioka, Toshihide Yamashita, David K. Crossman, Minae Niwa, Shin-ichi Kano

## Abstract

Various states of microglia appear in neuroinflammation, but their impact on brain function and behavior is not fully understood. Here we report that synapse engulfing microglia in the thalamus are crucial for cognitive impairment after cortical brain injury. Region-specific manipulations of reactive microglia in the chronic phase of injuries showed that microglial changes in the thalamus, but not in the hippocampus, impaired recognition memory. Single-cell RNA-sequencing analysis revealed the enrichment of synapse engulfing microglia in the thalamus, which developed in a CD9-dependent manner and caused synaptic loss and recognition memory deficits. In the thalamus, the blood-brain barrier was disrupted, and extravasated γ-immunoglobulins (IgG) co-localized with synapse engulfing microglia. Fc_γ_ receptor III blockade in the thalamus reduced synapse engulfing microglia, synapse loss, and recognition memory deficits. These findings demonstrate that the induction of synapse engulfing microglia in the thalamus by extravasated IgG/Fc_γ_RIII and CD9 signals causes recognition memory deficits after cortical brain injury.

## Introduction

Microglia are resident immune cells in the brain and play both neuroprotective and neurotoxic roles in neuroinflammation (*1–4*). Microglia react to infection and tissue destruction by recognizing danger molecules, secreting various inflammatory mediators, performing phagocytosis, and facilitating the infiltration of other immune cells. Reactive microglia also lose their homeostatic function to support neurons, consequently dysregulating neuronal activities and damaging neurons. Recent studies have revealed diverse states of reactive microglia under various pathological conditions, such as disease-associated microglia (DAM) - first described in mouse models carrying Alzheimer’s disease-related mutations (*5–7*). However, the impact of these microglial states on neurons and cognitive functions is not fully understood.

Traumatic brain injury (TBI) causes a long-lasting impact on the brain beyond acute injury, later inducing cognitive impairment and dementia (*8–13*). In the acute phase after TBI, reactive microglia try to maintain tissue integrity in the primary injury area; they change morphologies, produce neurotrophic factors and anti-inflammatory cytokines, and remove cellular debris via phagocytosis (*14–16*). In the chronic phase, reactive microglial changes are observed in multiple secondary injury areas beyond the primary injury area, including the hippocampus, thalamus, corpus callosum, putamen, supramarginal gyrus, and amygdala, and are associated with cognitive impairment (*14, 17, 18*). Nevertheless, the molecular mechanisms underlying microglial changes during secondary injuries and their impact on neuronal function remain elusive. Understanding such mechanisms would advance our understanding of the long-lasting impact of TBI on cognition and help identify effective therapeutic targets.

In this study, we have employed region-specific microglia depletion and manipulation to determine their roles in neuronal dysfunction and cognitive impairment in the chronic phase of cortical brain injuries. We have discovered that reactive microglia in the thalamus primarily drive recognition memory deficits after cortical brain injuries. Moreover, we have identified and inhibited the CD9-dependent generation of a distinct microglial state involved in synapse phagocytosis in the thalamus, showing its crucial role in thalamic synapse loss and recognition memory deficits. We have further elucidated that the blood-brain barrier (BBB) disruption and subsequent IgG extravasation in the thalamus induce synapse engulfing microglia via Fc receptor signaling, which cause thalamic synapse loss and cognitive impairment in the injured mice.

## Results

### Cortical brain injuries induced delayed thalamic microglia changes that correlate with recognition memory deficits in mice

To identify the reactive microglial changes most critical for cognitive impairment after cortical injuries, we first systematically assessed microglial changes across various brain regions using cortical ablation injury targeting the primary sensory and motor cortex (*19–21*) (**fig. S1A-D**). Increased levels of Iba1 protein expression, which is associated with reactive microglial changes, were observed in the primary cortical lesion area immediately after injury and were gradually attenuated toward day 21 post-injury (**Fig. 1A, B** and **fig. S1C, D**). In contrast, Iba1 expression in other brain regions began to increase around 7 days after injury. By 21 days after injury, the thalamus stood out as one of the regions with the most remarkable Iba1 expression (**Fig. 1A, B** and **fig. S1B-D**). The increased Iba1 expression was also associated with microglial morphological changes of rounder somas with shortened processes (**Fig. 1C** and **fig. S1B**). In the thalamus, microglial changes began in the ventrolateral nucleus (VL) around 7 days after injury and spread to the anterodorsal nucleus (AD), anteroventral nucleus (AV), centromedian nucleus (CM), ventral posterolateral nucleus (VPL), and mediodorsal nucleus (MD) by 21 days after injury (**fig. S1D**). We also observed similar microglial reactive changes in two other mouse cortical brain injury models, open controlled cortical impact (CCI) and repeated closed CCI (**fig. S1E-G**). In addition to the drastic morphological changes, thalamic microglia expressed high levels of a pro-inflammatory cytokine, tumor necrosis factor-_α_ (TNF-_α_), and a lysosome-associated protein, CD68 (**Fig. 1C, D**). CD68^+^ lysosomes in the thalamic microglia of injured mice were associated with PSD95, a postsynaptic scaffold protein, indicating microglial phagocytosis of synapses (**Fig. 1E**). Indeed, overall PSD95 signals in the thalamus significantly decreased in mice with cortical brain injuries compared to sham mice, indicating synaptic loss (**Fig. 1F**). Significant neuronal loss was also observed in the thalamus on day 21 post-injury (**fig. S1H, I**). These findings suggest that thalamic microglia engulf synaptic components and cause synaptic loss and neuronal death in the thalamus of mice with cortical brain injuries.

**Figure 1.**
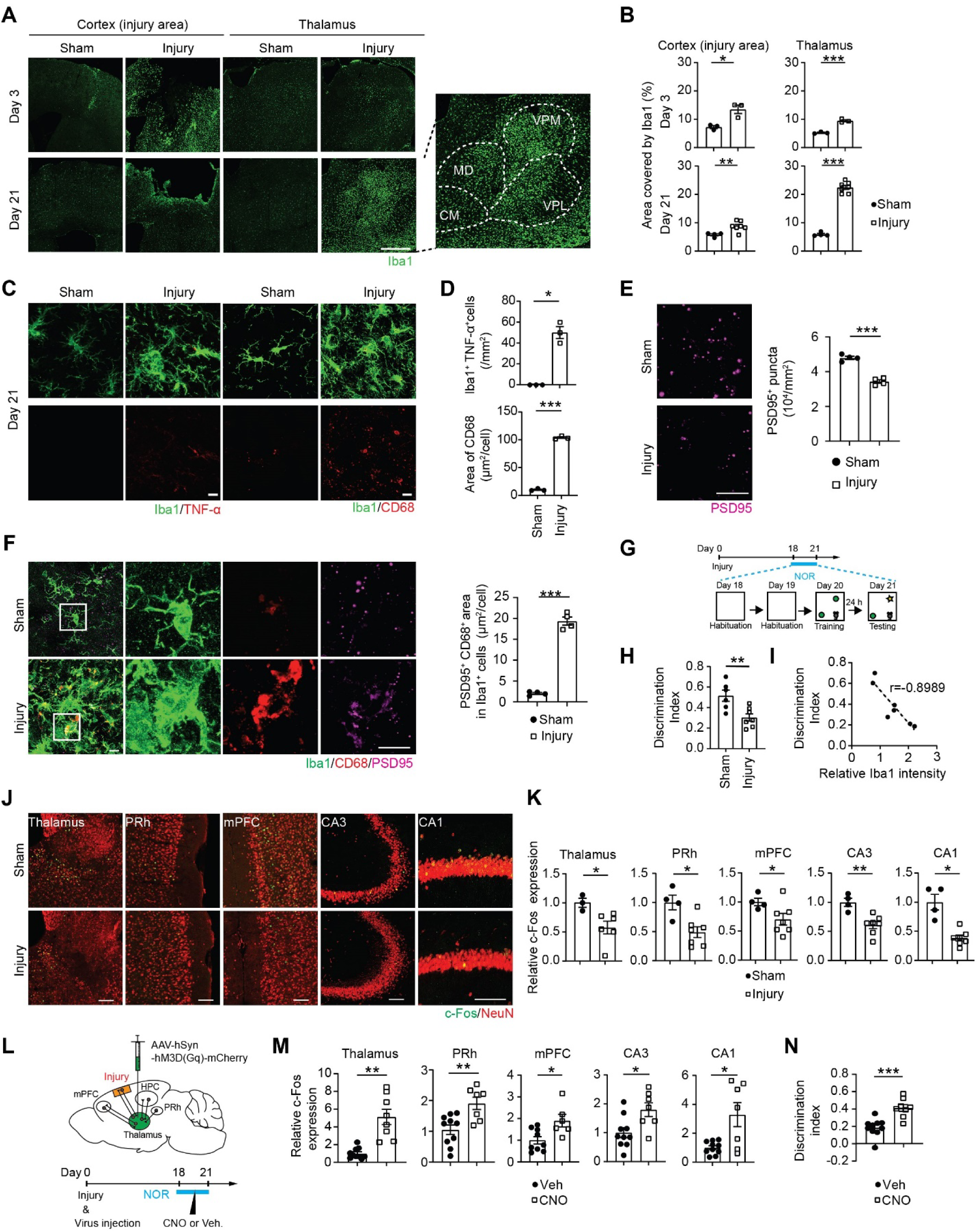
Cortical brain injuries induce reactive changes in the thalamic microglia, reduce thalamic neurons, and impair recognition memory deficits. (A) Representative images of Iba1^+^ microglia in the cortex (primary lesion) and the thalamus 3 and 21 days after unilateral cortical ablation injury and sham surgery. MD, mediodorsal nucleus; CM, central medial nucleus; VPL, ventral posterolateral nucleus; VPM, Ventral posteromedial nucleus. Scale bar, 500 µm. (B) Quantification data of the area covered by Iba1^+^ microglia (%) in the cortex and thalamus (n = 3-4 mice for sham; n = 3-7 mice for injury). (C) Representative images of TNF-α and CD68 expression in Iba1^+^ microglia in the thalamus 21 days after cortical brain injuries and sham surgery. Scale bar, 10 µm. (D) Quantification data of the number of TNF-α expressing Iba1^+^ microglia and average CD68^+^ area in Iba1^+^ microglia (n = 3 mice for sham; n = 3 mice for injury). (E) Representative images of PSD95^+^ signals overlapped with CD68^+^ area in Iba1^+^ microglia of the thalamus after cortical brain injuries and sham surgery. Scale bar: 10 µm. PSD95^+^ signals colocalized to CD68^+^ lysosomal area per Iba1^+^ microglia (PSD95^+^ CD68^+^ area inside Iba1^+^ microglia) were calculated per image, averaged across multiple brain sections per mouse and shown in bar graphs for the sham (n = 4 mice) and the injury (n = 4 mice) group. (F) Representative images of PSD95 post-synaptic proteins in the thalamus after cortical brain injuries and sham surgery. Scale bar: 10 µm. PSD95^+^ puncta density was calculated in the thalamus after cortical brain injuries and sham surgery. (n = 4 mice for sham; n = 4 mice for injury). (G) Experimental timeline for novel object recognition (NOR) test after cortical brain injuries. (H) Discrimination index in the NOR test for injury (n = 6) and sham (n = 7 mice) group. The discrimination index was calculated as ([time spent at the novel object] – [time spent at the familiar object])/([time spent at the novel object] + [time spent at the familiar object]). (I) Correlation plot of the discrimination index and thalamic Iba1 intensity 21 days after injuries and sham surgery (n = 7 mice). (J) Representative images of neuronal c-Fos expression in the thalamus, PRh, mPFC, and hippocampus (CA1, CA3) 21 days after injuries and sham surgery. Scale bars, 100 µm. (K) Quantification data of neuronal c-Fos expression in the injury group (n = 6-7 mice) relative to the sham group (n = 3-4 mice). (L) Experimental timeline for neuronal DREADD rescue experiment. AAV-hSyn-hM3D (Gq)-mCherry was injected into the thalamus on the day of cortical brain injuries, and CNO or vehicle was administered intraperitoneally before the test session in the NOR test. (M) Quantification data of neuronal c-Fos expression in the CNO-treated group (n = 8 mice) relative to the vehicle-treated mice (n = 11 mice). (N) Discrimination index in the NOR test for vehicle-treated (n = 11 mice) and CNO-treated mice (n = 8 mice). In bar graphs, data are shown as the mean□±□s.e.m, and each dot represents an individual animal. \**p* < 0.05, \*\**p* < 0.01, \*\*\**p* < 0.001; Student’s *t*-test (B, D, F, K, M, and N) and one-way ANOVA with *post hoc* Dunnett’s test (I). See also Figures S1-S3.

Cortical brain injuries also led to deficits in the novel objective recognition (NOR) test on day 21 post-injury (**Fig. 1G, H)**. Notably, a strong correlation was observed between NOR deficits (measured by discrimination index) and thalamic Iba1 staining intensity (**Fig. 1I**). The data is reminiscent of previous findings in human PET imaging studies where inflammatory signals related to microglia in the thalamus were correlated with cognitive impairment in TBI patients in a chronic phase of injuries (*17*). Although we primarily used female mice in this study, reactive microglial changes and NOR deficits after cortical injuries were similarly observed in male mice (**fig. S2A, B**). Collectively, our data indicate that thalamic microglia changes may be a critical step leading to cognitive impairment after cortical brain injuries.

### Thalamic neuronal dysfunction triggered cognitive impairment after cortical brain injuries

Reduced expression of neuronal c-Fos proteins, an indicator of neuronal activity, was also observed in the thalamus of injured mice after the NOR test, indicating an impaired activation of thalamic neurons (**Fig. 1J, K**). The NOR behavior is regulated by a network of neurons in multiple brain regions, including the perirhinal cortex (PRh), medial prefrontal cortex (mPFC), and hippocampus (HPC), in addition to the thalamus (*22, 23*). Notably, reduced c-Fos expression was also observed in the PRh, mPFC, and HPC. These results indicated that thalamic neuronal dysfunction could cause recognition memory deficits by negatively impacting the other neuronal activities linked to NOR. To address this possibility, we locally enhanced thalamic neuronal activities using a chemogenetic approach with the Designer Receptors Exclusively Activated by Designer Drugs (DREADD) in the injured mice (**Fig. 1L**) (*24*). Adeno-associated virus vectors expressing an activator DREADD [AAV-hSyn-hM3D(Gq)-mCherry] were injected into the thalamus of mice that received cortical injuries. Then, clozapine *N*-oxide (CNO) was administered intraperitoneally to activate DREADD. As expected, neuronal c-Fos expression increased in the thalamus of injured mice upon CNO administration (**Fig. 1M** and **fig. S3A**). Neuronal c-Fos expression also increased in the PRh, mPFC, and HPC, supporting our hypothesis that reduced thalamic neuronal activities impact neuronal activities in these brain regions. Under this condition, the CNO-treated injured mice showed an improved NOR (**Fig. 1N**). CNO injection alone did not affect microglial reactive changes, neuronal c-Fos expression, or NOR in the injured mice (**fig. S3B-D**). Thus, our data demonstrate that the reduced neuronal activities in the thalamus after cortical injuries trigger NOR deficits.

### Microglial reactive changes in the thalamus caused neuronal dysfunction and cognitive impairment after cortical injuries

To address the role of thalamic microglial reactive changes in neuronal dysfunction and cognitive impairment, we locally injected anti-colony stimulating factor 1 receptor antibodies (anti-CSF1R Ab), which have been used to deplete macrophages and microglia (*25–28*), into the thalamus of injured mice via implanted cannulas (**Fig. 2A**). As expected, intra-thalamic anti-CSF1R Ab injection reduced the number of reactive microglia in the thalamus (**fig. S4A**). Notably, anti-CSF1R Ab injection into the injured mice attenuated their NOR deficits, restored neuronal c-Fos expression in the thalamus and other areas related to NOR, and reduced neuronal loss in the thalamus (**Fig. 2B-D** and **fig. S4B**). Anti-CSF1R Ab injection also reduced microglia and restored NOR in the mice after the open CCI (**fig. S4C, D**). Hippocampal (HPC) microglia have been widely investigated in cognitive impairment after cortical brain injuries (*29, 30*). Unexpectedly, local injection of anti-CSF1R Ab into the HPC of injured mice did not restore NOR (**Fig. 2E, F** and **fig. S4E**). Indeed, microglial changes in the HPC of injured mice were more subtle than those in the thalamus, with little expression of translocator protein (TSPO), a mitochondrial activation marker and an established target of microglia tracer in human PET imaging (*17, 31–33*) (**Fig. 2G, H**). Neuronal loss was also not evident in the HPC (**fig. S4F**). These data show that reactive microglial changes after cortical brain injuries vary across brain regions, and thalamic microglial changes are required for injury-induced NOR deficits.

**Figure 2.**
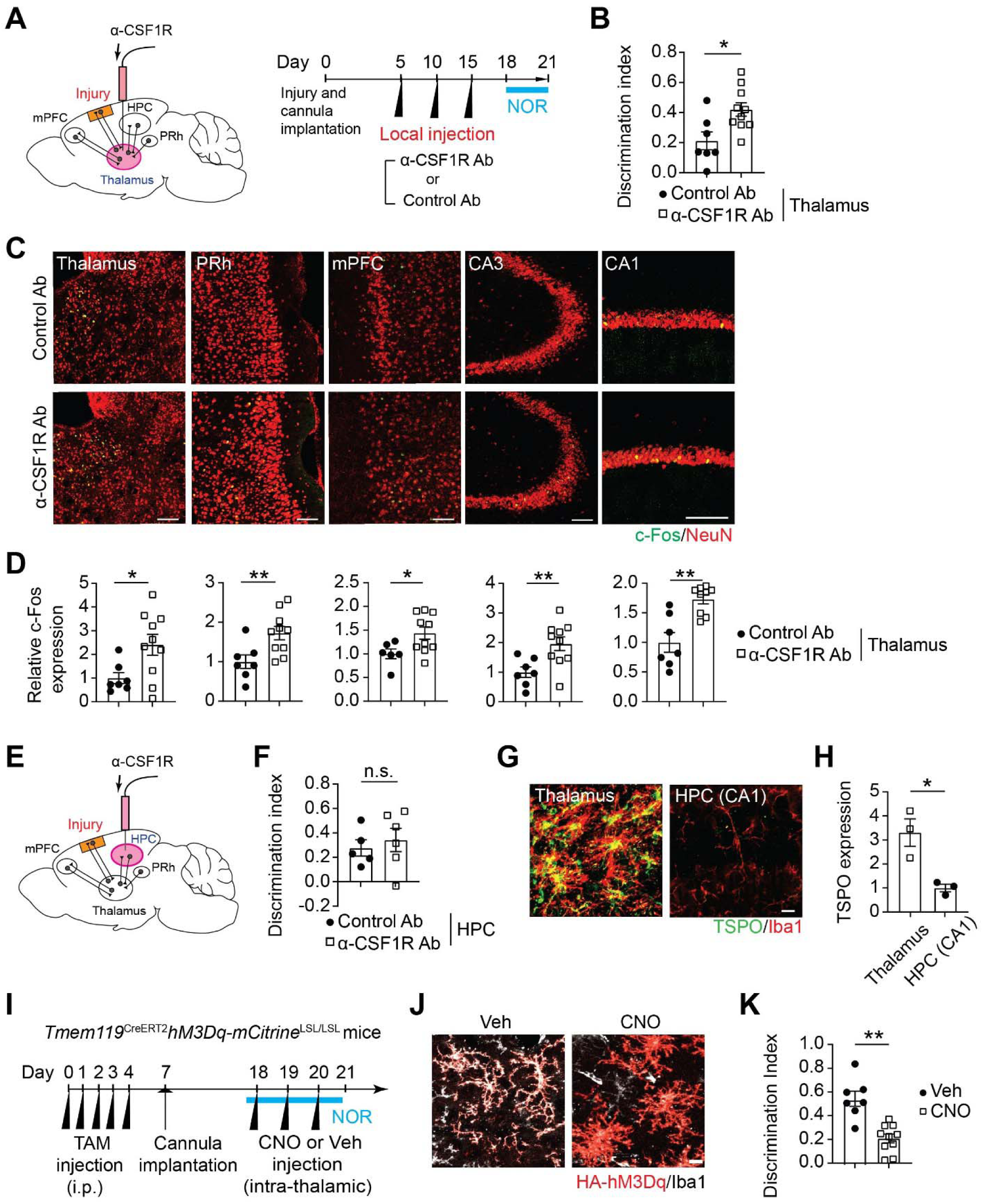
Reactive changes in thalamic microglia are necessary and sufficient to impair neuronal function and recognition memory. (A) Experimental timeline for local depletion of thalamic microglia by anti-CSF1R Ab injection. (B) Discrimination index in the NOR test for control Ab group (n = 7 mice) and anti-CSF1R Ab group (n = 10 mice) after cortical brain injuries. (C) Representative images of neuronal c-Fos expression in the thalamus, PRh, mPFC, and hippocampus (CA1, CA3) in control Ab- and anti-CSF1R Ab-injected group after cortical brain injuries. Scale bars, 100 µm. (D) Quantification data of neuronal c-Fos expression in the anti-CSF1R Ab group (n = 10 mice) relative to the control Ab group (n = 7 mice). (E) Experimental strategy for local depletion of hippocampus microglia by anti-CSF1R Ab injection. (F) Discrimination index in the NOR test for the mice with intrahippocampal injections of control Ab (n = 5 mice) and anti-CSF1R Ab (n = 6 mice). (G) Representative images of TSPO expression in Iba1^+^ microglia of the thalamus and HPC (CA1) 21 days after cortical brain injury. Scale bar, 10 µm. (H) Quantification data of microglial TSPO expression in HPC relative to thalamus (n = 3 mice per region). (I) Experimental timeline for DREADD experiments. *Tmem119 ^CreERT2^hM3Dq-mCitrine^LSL/LSL^*mice received intraperitoneal (i.p.) injections of tamoxifen (TAM) injections for five consecutive days to induce hM3D(Gq)-mCitrine expression in Tmem119^+^ microglia. Then, the mice were implanted with cannulas into the thalamus. Two weeks after the cannula implantation, the mice underwent the NOR test. CNO or vehicle (Veh) was i.p. injected 30 min before the habituation and training sessions of the NOR test. (J) Representative images of HA-tagged hM3D(Gq) DREADD expression in Iba1^+^ microglia in the thalamus in *Tmem119 ^CreERT2^hM3Dq-mCitrine^LSL/LSL^* mice after TAM injections and CNO or vehicle (Veh) injection. Scale bar, 100 µm. (K) Discrimination index in the NOR test for vehicle (Veh)- (n = 7 mice) and CNO-treated group (n = 10 mice). In bar graphs, data are shown as the mean□±□s.e.m, and each dot represents an individual animal. \**p* < 0.05, \*\**p* < 0.01; n.s., not significant; Student’s *t*-test.

We next tested whether reactive changes of thalamic microglia were sufficient to impair NOR in non-injured mice. We first injected lipopolysaccharides (LPS) locally into the thalamus of non-injured mice via cannulas to induce microglial reactive changes (**fig. S5A**). LPS injection resulted in remarkable thalamic microglial reactive changes and NOR deficits, accompanied by reduced neuronal c-Fos expression in the thalamus and other NOR-related brain regions (**fig. S5B-D**). We also used a recently reported method of chemogenetic activation of microglia (*34*). We generated a conditional transgenic mouse line in which an activator DREADD was expressed in microglia (*Tmem119 ^CreERT2^hM3Dq-mCitrine^LSL/LSL^*mice) and induced thalamic microglial reactive changes by local CNO injection into the thalamus (**Fig. 2I**). DREADD activation caused reactive microglial changes locally in the thalamus, and impaired NOR and neuronal c-Fos expression (**Fig. 2J, K** and **fig. S6A-D**). These data demonstrate that reactive microglial changes in the thalamus are sufficient to cause NOR deficits and related neuronal dysfunction similar to those in cortically injured mice.

### A distinct microglial state with high expression of *Cd9* and phagocytosis-related genes increased in the thalamus of injured mice

To better understand the molecular mechanisms underlying thalamic microglial changes in injured mice, we performed single-cell RNA sequencing (scRNA-seq) of CD45^+^ cells in the thalamus and HPC of injured mice. We identified 19 transcriptionally distinct cell types and states, with 6 distinct states belonging to microglia (designated as MG#1-6) (**Fig 3A**, **fig. S7A**, and **Table S1**). The 6 distinct microglial states were then compared to previously reported microglial states, including homeostatic microglia (HM), disease-associated microglia (DAM), and interferon-responsive microglia (IRM) (**Fig. 3B, C**, **fig. S7B**, and **Table S2**)(*35*). MG#2 and #3 were enriched with HM-related genes, such as *P2ry12* and *Tmem11*9. MG#4 was enriched with DAM-related genes, such as *Cd9* and *Lpl*. MG#6 was enriched with IRM-related genes, such as *Ifitm3* and *Stat1*. MG#1 had an intermediate state that partially overlapped with HM, DAM, and IRM. MG#5 was different from the other states, showing a significantly reduced expression of ribosomal protein genes, such as *Rps16*, *Rps23*, and *Rpl19*. Pseudotime trajectory analysis revealed that MG#2 and #3 progressed toward MG#4, #5, or #6 through several common intermediate states in MG#1 (**Fig. 3D** and **Table S3**).

**Figure 3.**
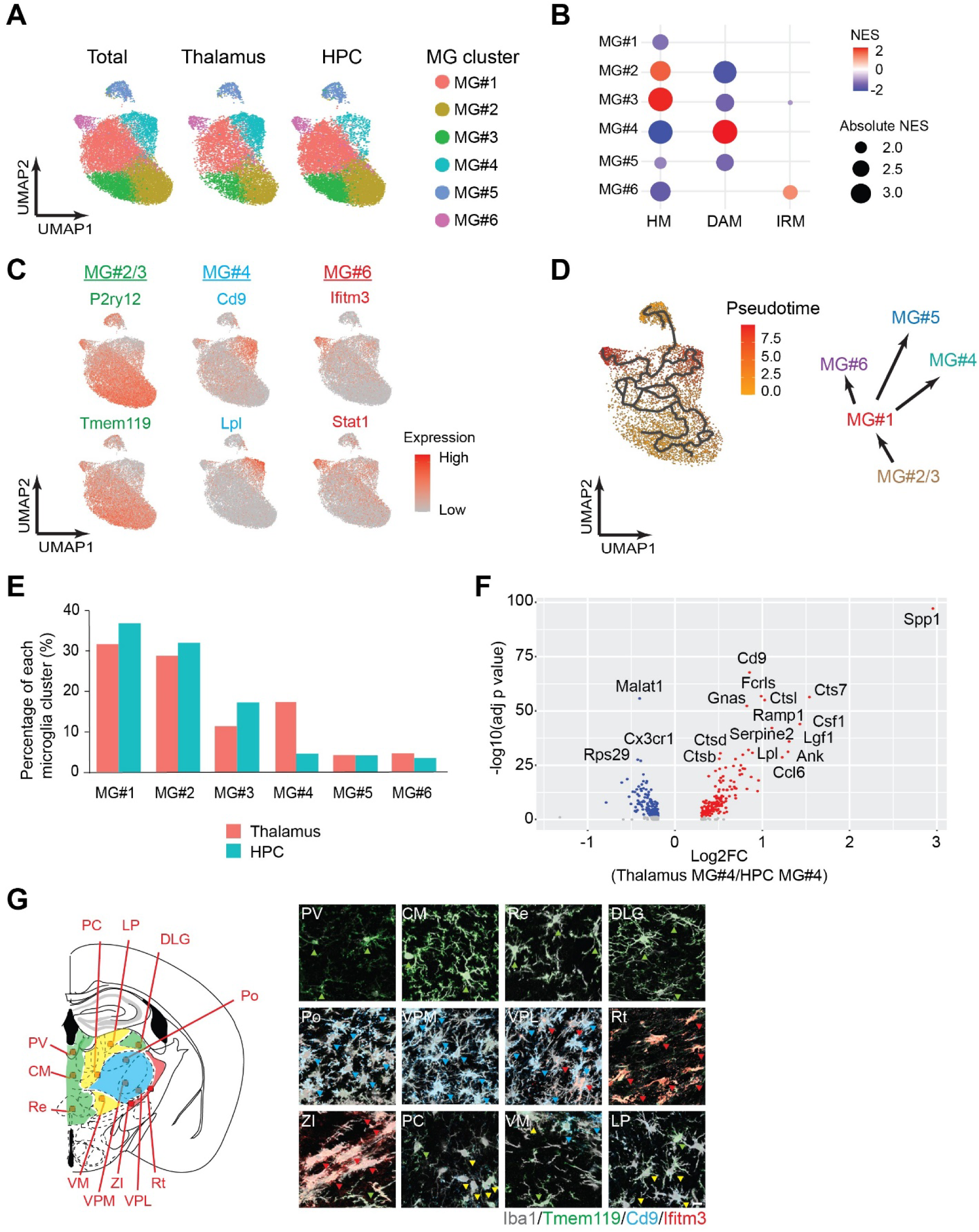
Microglia with high phagocytic capability are enriched in the thalamic inflammation secondary to cortical injury. (A) Uniform manifold approximation and projection (UMAP) plot highlighting 6 microglia clusters after cortical brain injuries. (B) Dot plots showing enrichment of gene expression features for HM, DAM, and IRM across 6 microglia clusters based on the gene set enrichment analysis (GSEA). The circle sizes reflect absolute NES, and their colors indicated positive (red) and negative (blue) values. The data were corrected for multiple comparisons with the Benjamini-Hochberg method, and only the significant results (at adjusted p-values below 0.05) were shown. (C) Feature plots showing the genes enriched in MG#2/3 (*P2ry12*, *Tmem119*), MG#4 (*Cd9, Lpl*), MG#6 (*Ifitm3*, and *Stat1*) across all the microglia clusters. (D) Pseudo-time analysis showing a cell transition trajectory from MG#2/3 to MG#4, #5, or #6 via MG#1 and a schematic diagram showing how MG#2 and #3 transit to MG#4, #5, and #6 via MG#1. (E) Bar graphs showing the percentages of each microglial cluster in the thalamus (red) and hippocampus (blue), with the percentages reaching 100% for each brain region. (F) Volcano plot of differentially expressed genes in the thalamic MG#4 compared to the HPC MG#4. Red: genes upregulated in thalamic MG#4. Blue: genes downregulated in thalamic MG#4. (G) Spatial distribution of MG#1, MG#2/3, MG#4, and MG#6 in the thalamus 21 days after cortical injuries. Brain areas predominantly occupied by microglia with distinct states are shown with respective colors: MG#1 (yellow), MG#2/3 (green), MG#4 (blue), and MG#6 (red). Representative images of MG#1 (Tmem119^lo^ CD9^lo^ Iba1^+^, yellow arrowheads), MG#2/3 (Tmem119^hi^ Iba1^+^, green arrowheads), MG#4 (CD9^hi^ Iba1^+^, blue arrowheads), and MG#6 (Ifitm3^hi^ Iba1^+^, red arrowheads) are shown in each thalamic subregion: paravetricular nucleus (PV), central medial nucleus (CM), nucleus reuniens (Re), dorsal lateral geniculate nucleus (DLG), posterior nucleus (Po), ventral posteomedial nucleus (VPM), ventral posteorateral nucleus (VPL), reticular nucleus (Rt), zona incerta (ZI), paracentral nucleus (PC), ventromedial nucleus (VM), and lateral posterior nucleus (LP). Scale bar: 10 µm.

Although MG#4 showed an overlap with DAM in their gene expression profiles, the expression of *Trem2*, a representative gene of DAM (*36–38*), was not restricted to MG#4 (**fig. S7C**). We also observed that microglia with low CD9 and high Trem2 expression (CD9^lo^Trem2^hi^ microglia) were present both in the thalamus and the HPC, whereas microglia with high CD9 expression (both CD9^hi^ Trem2^lo^ and CD9^hi^ Trem2^hi^ microglia) were enriched in the thelamus (**fig. S7D, E**). These data suggested that MG#4 might be a state different from Trem2 expressing DAM. Notably, MG#4 was more than three-fold enriched in the thalamus than in the HPC (**Fig. 3E**). In addition, thalamic MG#4 is distinct from HPC MG#4; genes related to phagocytosis (*Spp1*, *Cd9,* and *Lpl)* and lysosomes (*Cts7, Ctsl, Ctsb,* and *Ctsd*) were significantly higher in the thalamic MG#4 than the HPC MG#4 (**Fig. 3F** and **Table S4**). Immunohistochemistry data confirmed a substantially higher number of CD9^hi^ Iba1^+^ cells in the thalamus than the HPC (**fig. S7F**). Thalamic CD9^hi^ Iba1^+^ cells also co-expressed LPL proteins (**fig. S7G**). These findings indicated MG#4 in the thalamus had enhanced phagocytotic activities compared to those in the HPC.

We next examined the spatial distribution of microglia with distinct states in the thalamus of injured mice by immunohistochemistry (Fig. 3G). MG#2 and #3, visualized as Tmem119 Iba1 cells, were found in a region distant from the center of the accumulation of microglia, such as the paraventricular nucleus (PV), central medial nucleus (CM), nucleus reuniens (Re), and dorsal lateral geniculate nucleus (DLG). MG#4, visualized as CD9^hi^ Iba1^+^ cells, were distributed around the posterior nucleus (Po), ventral posteromedial nucleus (VPM), and ventral posterolateral nucleus (VPL), regions where microglial morphological changes were most remarkable. MG#6, visualized as Ifitm3^hi^ Iba1^+^ cells, was found in the reticular nucleus (Rt) and zona incerta (ZI). MG#1, visualized as cells expressing Tmem119 and CD9 at lower levels, was observed in the paracentral nucleus (PC), ventromedial nucleus (VM), and lateral posterior nucleus (LP). In summary, MG#4 (CD9^hi^) was highly accumulated in the area where the most remarkable microglial morphological changes and the highest expression of Iba1 were observed (blue area); MG#6 (Iftm3^hi^) was located in the periphery (pink area); MG#2 and #3 (Tmem119^hi^) were located at distant sites (green area); and MG#1 (Tmem119^lo^CD9^lo^) was located between the areas occupied by MG#4 and #2/3 (yellow area) (**Fig. 3G**). The spatial localization of MG#4 in the inflammation core indicates their pivotal role in thalamic inflammation and neuronal dysfunction after cortical injury.

### Synapse engulfing thalamic microglia developed from reactive microglia precursors in a CD9-dependent manner and drove recognition memory deficits

We then tested if MG#4 caused synaptic loss by phagocytosis in the thalamus of injured mice, imparing recognition memory. As MG#4 highly expressed CD9, we reasoned that CD9 blockade could inhibit the generation of MG#4. Previous studies used anti-CD9 blocking Ab to inhibit macrophage phagocytosis and other biological processes under *in vitro* and *in vivo* conditions (*39–41*). Therefore, we locally injected anti-CD9 blocking Ab into the thalamus via cannulas to assess the impact of MG#4 inhibition on thalamic pathology and recognition memory deficits (**Fig. 4A**). Multiple anti-CD9 Ab local injections from 7 to 19 days post-injury reduced the number of CD9^hi^ microglia, the area covered by CD68^+^ lysosomes per microglia and the amount of PSD95 in CD68^+^ area per microglia on day 21 post-injury (**Fig. 4B** and **fig. S8A, B**), indicating the attenuated microglial reactivities and phagocytosis of synapses. Accordingly, PSD95^+^ synaptic puncta, neuron numbers, and neuronal c-Fos expression increased in the thalamus of anti-CD9-injected injured mice (**fig. S8C-E**). Consistent with these histological findings, CD9 blockade in the thalamus attenuated NOR deficits of injured mice (**Fig. 4C**). These data show that MG#4 (CD9^hi^ microglia) are critical for thalamic microglial synapse phagocytosis, synapse loss, and recognition memory deficits following cortical brain injury.

**Figure 4.**
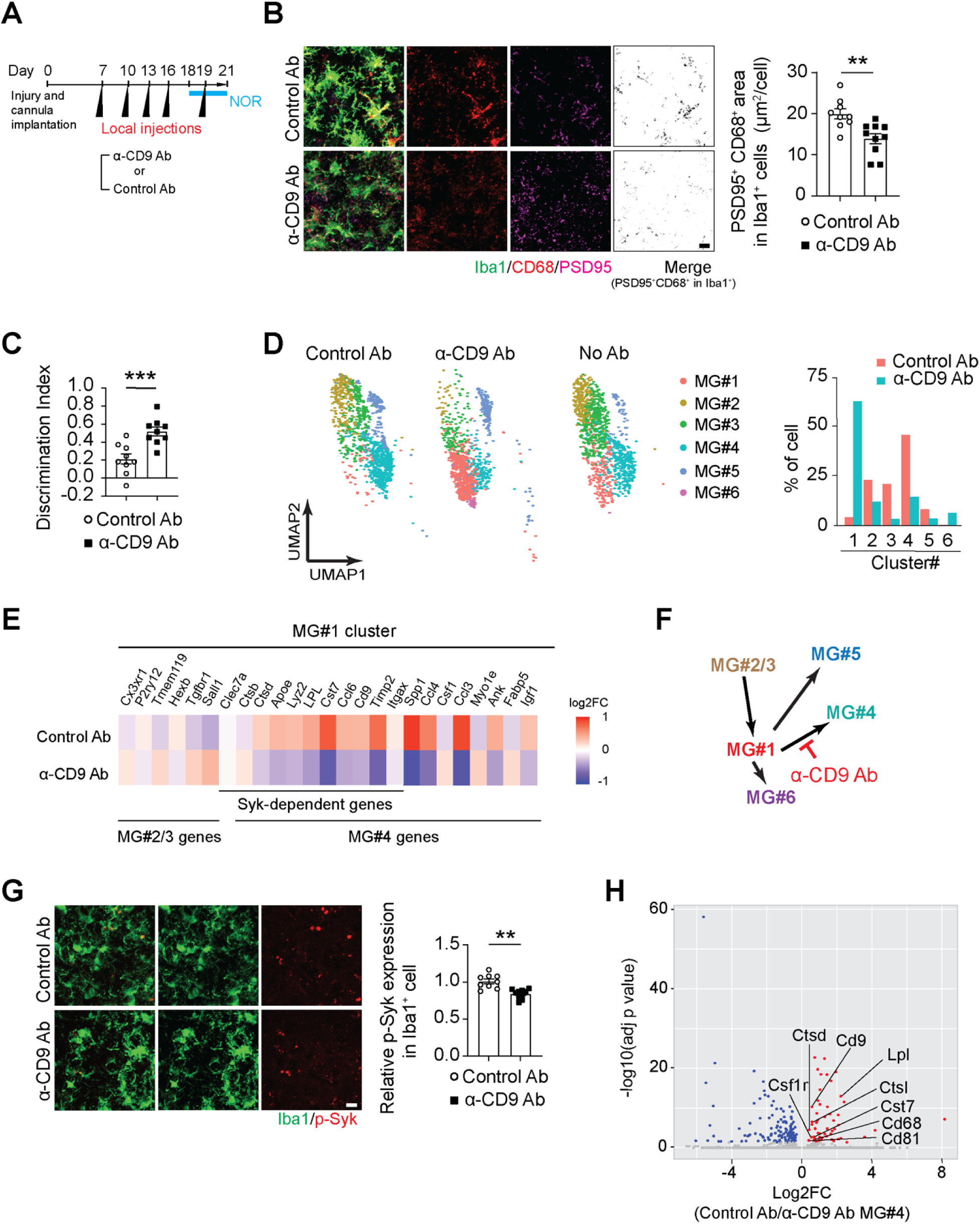
Local CD9 blockade in the thalamus inhibits the generation of synapse engulfing microglia from intermediate microglia, reduces synaptic loss, and attenuates cognitive impairment after cortical injuries. (A) Experimental timeline for blocking CD9 by anti-CD9 Ab injection. (B) Representative images of PSD95^+^ signals overlapped with CD68^+^ lysosomal area of Iba1^+^ microglia in the thalamus of control Ab- and anti-CD9 Ab-injected mice after cortical brain injuries. PSD95^+^ puncta colocalized to CD68^+^ lysosomal area in Iba1^+^ microglia (PSD95^+^ CD68^+^ Iba1^+^ area) are calculated per animal and shown in bar graphs for the control Ab- (n= 9 mice) and the anti-CD9 Ab-injected group (n = 10 mice). Scale bar: 10 µm. (C) Discrimination index in the NOR test for the control Ab (n = 9 mice) and the anti-CD9 Ab group (n = 10 mice). (D) UMAP plot highlighting 6 microglia clusters after cortical brain injuries in control Ab-injected group thalamus, in anti-CD9 Ab-injected group thalamus, and in thalamus from Fig.3A, and bar graphs showing the percentages of each microglial cluster in control Ab group (red) and in anti-CD9 Ab group (blue), with the percentages reaching 100% for each group. (E) Pseudo-bulk heat map showing log2FC of mean gene expression in thalamic MG#1 between the anti-CD9 Ab group and control Ab group, highlighting up- or down-regulated genes that are Syk-dependent or enriched in MG#2/3, MG#4, and MG#6. Positive values indicate higher expression (red) and negative values indicate lower expression in (blue). (F) A schematic diagram showing how MG#2 and #3 transit to MG#4, #5, and #6 via MG#1, and how anti-CD9 Ab acts on this pathway. (G) Representative images of p-Syk signals in Iba1^+^ microglia of the thalamus in control Ab- and anti-CD9 Ab-injected group after cortical brain injuries. Quantification data of the p-Syk levels in Iba1^+^ microglia in the anti-CD9 Ab group (n = 10 mice) relative to control Ab group (n = 9 mice) are shown in bar graphs. Scale bar: 10 µm. (H) Volcano plot of differentially expressed genes in the thalamic MG#4 microglia in anti-CD9 Ab group compared to the thalamic MG#4 microglia in control Ab group. Red: genes downregulated in anti-CD9 Ab group. Blue: genes upregulated in anti-CD9 Ab group. In bar graphs, data are shown as the mean□±□s.e.m, and each dot represents an individual animal. \*\**p* < 0.01, \*\*\**p* < 0.001; Student’s *t*-test.

To determine how CD9 blockade affects MG#4, we used scRNA-seq to examine thalamic microglial states in the injured mice receiving anti-CD9 Ab and control Ab. Six microglial subclusters appeared in the antibody-injected mice, similar to the injured mice without antibody injections (**Fig. 4D** and **fig. S8F, G)**. Anti-CD9 Ab injection significantly reduced MG#4 while increasing MG#1 (**Fig. 4D**) and reduced the expression of MG#4-related genes in MG#1 (**Fig. 4E** and **Table S5**), suggesting that anti-CD9 blocked the generation of MG#4 from MG#1 (**Fig. 4F**). CD9 blockade also reduced the phosphorylation of spleen tyrosine kinase (Syk) (**Fig. 4G**), which was known to mediate CD9 signaling (*42, 43*), in the thalamic microglia, and Syk-dependent gene expression signatures, overlapped with MG#4-related signatures, in MG#1 (**Fig. 4E**). Although we observed some MG#4 remained in the mice injected with anti-CD9 Ab, the expression of genes related to phagocytosis and lysosome was significantly lower in these MG#4 (**Fig. 4H** and **Table S6**). These results demonstrate that CD9 signaling is critical for activating the transcriptional program to develop MG#4 from MG#1 and for MG#4 to enhance synapse phagocytosis capability.

### Blood-brain barrier (BBB) disruption and IgG extravasation in the thalamus facilitated the generation of synapse engulfing microglia via Fc**_γ_** receptor III signaling

Our histological analysis further revealed that extravasated γ-immunoglobulins (IgG) significantly accumulated in the thalamus but not in the HPC of injured mice (**Fig. 5A** and **fig. S9A**). In the thalamus, the expression of Claudin5, a tight junction protein, significantly decreased compared to the HPC (**Fig. 5B**). Notably, anti-CD9 injections did not reduce IgG extravasation (**Fig. 5C**). Thus, BBB disruption and subsequent IgG extravasation preceded the generation of synapse engulfing thalamic microglia. CD9 was reported to be functionally associated with FcγRIII signaling in macrophages (*42*). Indeed, FcγRIII was expressed in CD9^hi^ microglia (**fig. S9B**). Thus, we tested if FcγRIII blockade affected the generation of synapse engulfing microglia in the thalamus (**Fig. 5D**). Local anti-FcγRIII Ab injections into the thalamus successfully reduced the number of CD9^hi^ microglia, attenuated microglial phagocytosis of synaptic component PSD95, and enhanced NOR in the injured mice (**Fig. 5E-H**). These findings suggest that BBB disruption and IgG extravasation in the thalamus facilitate the generation of synapse engulfing microglia (MG#4, CD9^hi^ microglia) via FcγRIII signaling, resulting in synaptic loss, neuronal dysfunction, and recognition memory deficits (**Fig. 5I**).

**Figure 5.**
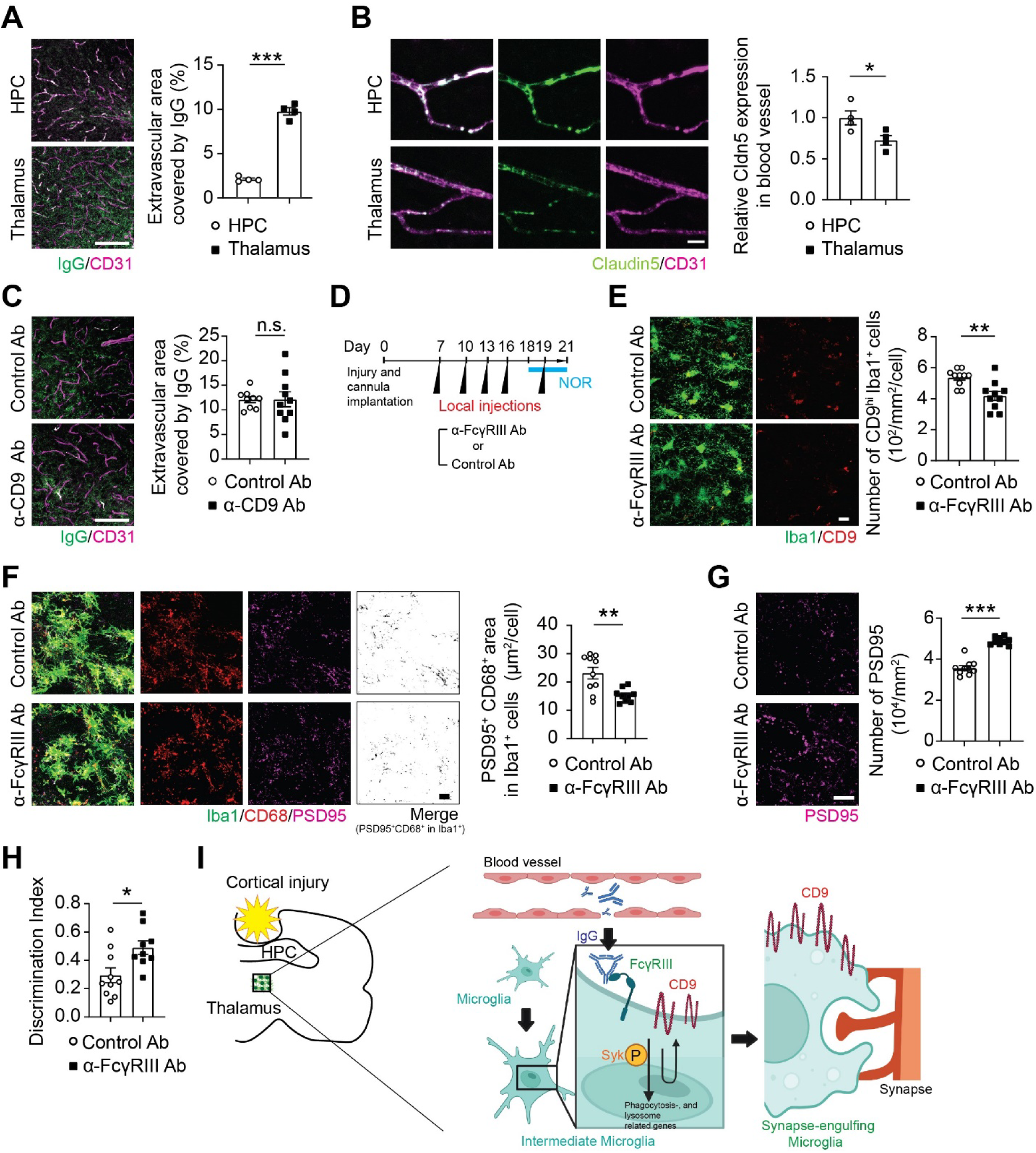
Extravasated IgGs induce the generation of synapse engulfing thalamic microglia via FcγRIII signaling. (A) Representative images of IgG and CD31^+^ blood vessels of the thalamus and the HPC after cortical brain injuries. Percentages of extravascular IgG coverage area per total IgG coverage area are shown in bar graphs for the HPC (n = 4 mice) and the thalamus (n = 4 mice) group. (B) Representative images of Claudin5 and CD31^+^ blood vessels in the hippocampus (HPC) and the thalamus after cortical brain injuries. Quantification of the relative expression of Claudin5 in CD31^+^ blood vessels in the HPC (n = 4 mice) and the thalamus (n = 4 mice) is shown in bar graphs. (C) Representative images of IgG and CD31^+^ blood vessels in the thalamus of cortically injured mice treated with control Ab and anti-CD9 Ab. Percentages of extravascular IgG coverage area per total IgG coverage area are shown in bar graphs for the control Ab (n = 10 mice) and the anti-CD9 Ab (n = 9 mice) group. (D) Experimental timeline for blocking Fcγ receptor III (FcγRIII) by anti-FcγRIII Ab injection. (E) Representative images of CD9^hi^ Iba1^+^ thalamic microglia in the cortically injured mice injected with control Ab and anti-FcγRIII Ab. Quantification data are shown in bar graphs for the control Ab (n = 10 mice) and the anti-FcγRIII Ab group (n = 9 mice). (F) Representative images of PSD95^+^ signals overlapped with CD68^+^ lysosomal area in Iba1^+^ microglia of the thalamus in control Ab and anti-FcγRIII Ab injected mice after cortical brain injuries. PSD95^+^ signals in CD68^+^ lysosomes per Iba1^+^microglia are calculated as PSD95^+^ CD68^+^ area and shown in bar graphs for control Ab (n= 10 mice) and anti-FcγRIII Ab group (n = 9 mice). (G) Representative images of PSD95^+^ synaptic puncta of the thalamus in control Ab- and anti-FcγRIII Ab-injected mice after cortical brain injuries. Quantification data are shown in bar graphs for control (n = 10 mice) and anti-CD9 Ab-injected group (n = 9 mice). (H) Discrimination index in the NOR for control Ab (n = 10 mice) and anti-FcγRIII Ab (n = 9 mice) group. (I) Graphic illustration showing how synapse-engulfing microglia develop in response to extravasated IgG via FcγRIII-CD9 signaling, engulfing synapses and impairing neuronal function. This illustration was created with help of BioRender.com. Scale bars, 10 µm (A, E, F, and G) and 100 µm (C). In bar graphs, data are shown as the mean□±□s.e.m, and each dot represents an individual animal. \**p* < 0.05; \*\**p* < 0.01; \*\*\**p* < 0.001; n.s., not significant; Student’s *t*-test.

## Discussion

In this study, we have demonstrated that reactive microglial changes in the thalamus during secondary injuries following cortical brain injuries are crucial for neuronal dysfunction and recognition memory deficits. We have discovered that synapse engulfing microglia (MG#4), expressing high levels of genes related to phagocytosis and lysosome function, are enriched in the thalamus in the chronic phase of cortical brain injury. These synapse engulfing microglia in the thalamus develop in a CD9-dependent manner and are responsible for synapse loss and recognition memory deficits. Our data have further shown that the BBB disruption and subsequent IgG extravasation facilitate the generation of synapse engulfing microglia via FcγRIII signaling. Thus, our study has identified a spatially and temporally restricted generation of a distinct state of synapse engulfing microglia that mediates the transition of acute cortical brain injury into chronic cognitive impairment.

Diverse microglial states have been defined by their gene expression signatures under various neuroinflammatory conditions (*5–7, 35*). Nevertheless, their impact on neuropathology and cognitive impairment has not been fully understood. This is partly because the established tools to investigate microglia *in vivo*, including pharmacological depletion (e.g., PLX compounds) (*44*) and Cre transgenic mice (e.g., *Cx3Cr1*^CreERT2^ mice), do not allow either region- or state-specific manipulation of microglia. As a result, the roles of microglia in brain pathology and dysfunction have been occasionally controversial. In rodent TBI models, some studies reported that the repopulation of fresh microglia after depletion attenuated cognitive impairment by secreting neurotrophic factors (*29, 45*), while others claimed that microglial depletion prevented the development of cognitive impairment (*46*). In this study, we have used region-specific microglia manipulations using intracranial injections of blocking antibodies via implanted cannula to address the causal role of reactive microglia in cognitive impairment following cortical brain injury. This approach has helped us identify a critical role of thalamic microglial changes in cortical injury-induced recognition memory deficits. As regional heterogeneity in microglia exists in various brain disorders, similar approaches may be helpful in understanding the relative contributions of region-specific microglia to brain pathology and dysfunction in other disease contexts.

Our scRNA-seq analysis has revealed the microglia with multiple distinct states in the two representative secondary injury areas (the thalamus and the HPC) in the chronic phase of cortical brain injury, including CD9^hi^ microglia (MG#4), Ifitm3^hi^ IFN-responsive microglia (MG#6), Tmem119^hi^ homeostatic microglia (MG#2, #3), and microglia with intermediate phenotype (MG#1). Although MG#4 is characterized by the enriched expression of DAM markers, such as *Cd9* and *Lpl,* the expression of *Trem2* is not selectively enriched in MG#4. MG#4 also causes synapse loss and impairs cognition, while DAM plays a neuroprotective role (*5*). These results suggest that MG#4 is functionally distinct from DAM. Intriguingly, these microglia show distinct spatial localization patterns within the thalamus depending on their states; synapse engulfing CD9^hi^ microglia (MG#4) and IFN-responsive microglia (MG#6) are localized to distinct thalamic subregions in an almost mutually exclusive manner. Several recent studies using scRNA-seq have also shown the presence of microglia with distinct transcriptional states in TBI (*47–50*). However, few studies have examined reactive microglial changes in the secondary injury areas. Thus, our scRNA-seq data have generated a novel molecular insight into the reactive microglial changes in the secondary injuries that are closely associated with cognitive impairment after cortical brain injury. Future studies would dissect the intimate relationships among these spatially localized distinct microglial states.

Mechanisms to induce synapse engulfing microglia are not fully understood. Our data demonstrate that CD9 signaling is crucial for the development of a distinct microglial state involved in synapse phagocytosis (MG#4) in the thalamus of cortically injured mice. Notably, Syk, a downstream kinase for CD9 signaling, has been recently shown to play a critical role in microglial phagocytosis of neurotoxic materials in mouse models of Alzheimer’s disease and multiple sclerosis (*51, 52*). As CD9 is reported to be expressed in several microglial states, such as DAM, under Alzheimer’s disease and other neurodegenerative conditions, CD9-dependent generation of phagocytic microglia may be a shared mechanism across multiple disease contexts. Our data also show that CD9 blockade inhibits the expression of a broad range of Syk-independent genes related to MG#4 in MG#1. Thus, CD9 signaling may have additional mediators other than Syk, which can be targeted to control microglial synapse phagocytosis. Previous studies report that TREM2 is required for reactive microglia to become DAM (*5*). Although MG#4 and DAM have an overlapped gene expression signature, our data reveal that CD9 expression is more confined to MG#4 while Trem2 is expressed across multiple reactive states in the thalamic microglia in the chronic phase of cortical brain injury. Thus, CD9 and Trem2 may have differential contributions to the generation of microglia with vigorous phagocytic activities. Both CD9 and Trem2 are cell surface molecules enriched in specific microglial states. Further studies focusing on the role of microglial state-specific cell surface molecules may help reveal the mechanisms underlying the transition of reactive microglial states to specific phenotypes, such as phagocytic microglia.

Our findings also reveal a previously unrecognized role of FcγRIII signaling in the induction of phagocytic phenotypes in microglia. Engagement of activating type I FcγR, such as FcγRIII, and stimulation of downstream signaling pathways by IgG has been shown to induce transcriptions of genes encoding pro-inflammatory mediators, phagocytosis, and lysosomes in neutrophils, macrophages, and other phagocytic cells (*53*). FcγR signaling is also known to play a critical role in generating osteoclasts, which are highly phagocytic cells with elevated expression of osteopontin (Spp1) and CD9 (*54–56*). Thus, FcγR signaling-mediated induction of phagocytic phenotypes may be shared features across multiple myeloid lineage cells.

This study has found that BBB disruption and IgG extravasation occur predominantly in the thalamus during the chronic phase of cortical brain injury. One possible explanation for this selectivity of BBB disruption in the thalamus is that damages to the highly dense connections between the thalamus and primary cortical injury areas result in the release of excessive axonal debris, inducing massive inflammation involving both microglia and astrocytes, the latter contributing to the BBB breakage. Another possibility is that microglia in the thalamus may be more prone to damage the BBB upon reactive changes than those in other brain areas. Regional differences in microglia, as recently reported (*57, 58*), may also underlie BBB disruption and IgG extravasation in the thalamus. Irrespective of the underlying mechanisms, targeting the BBB disruption and IgG/FcγRIII-dependent signaling in microglia may provide potential therapeutic strategies to prevent chronic neuroinflammation and cognitive impairment after cortical brain injury.

Human positron emission tomography (PET) imaging studies using a translocator protein (TSPO) ligand have consistently revealed persistent thalamic inflammation long after TBI, which is correlated with cognitive impairment (*17, 33*). Postmortem brain studies have also demonstrated that thalamic damage and neuroinflammation are prevalent in TBI and associated with worse outcomes (*59, 60*). Rodent studies have also observed significant microglial changes in the thalamus post-TBI (*47, 61*). Our findings align with these previous observations and further elucidate a pivotal role for thalamic microglial changes in the progression from acute brain injury to chronic cognitive impairment. Further studies on human TBI would reveal the detailed localization of microglia with distinct states in the brains after cortical injuries and advance our understanding of microglial contributions to cognitive impairment, contributing to the development of effective therapeutic interventions to prevent post-TBI cognitive impairment.

## Materials and Methods

### Study design

The main objective of this study was to elucidate the role of reactive microglial changes in cognitive impairment after cortical brain injuries using mouse models, focusing on thalamic microglia. We implanted a cannula in the brain and locally injected antibodies and chemical reagents to elucidate the roles of thalamic microglial reactive changes, CD9 and FcγRIII signals. Individual mice were randomly assigned to experimental groups. Investigators were not blinded during data acquisition and analysis. The number of independent experiments performed, n, is indicated in the figure legends.

### Mice

C57BL/6J (#664), Tmem119-CreERT2 (*Tmem119^CreERT2^*, #031820), CAG-LSL-hM3Dq-mCitrine (*hM3Dq-mCitrine^LSL/LSL^,* #026220), and B6.Cg-Tg (APP SwFlLon,PSEN1*M146L* L286V)6799Vas/Mmjax, #034848) mice were purchased from the Jackson Laboratory. *Tmem119^CreERT2^* and *hM3Dq-mCitrine^LSL/LSL^*mice were crossed to generate *Tmem119 ^CreERT2^hM3Dq-mCitrine^LSL/LSL^*mice. Female mice were used for most of the experiments unless stated otherwise. All the mice were housed in specific pathogen-free facilities with *ad libitum* access to food and water under a standard light/dark cycle at the University of Alabama at Birmingham. All experimental procedures were performed under the animal protocols approved by the Institutional Animal Care and Use Committees.

### Cortical brain injuries

#### Cortical Ablation Injury (CAI)

Unilateral lesions of the cortex were induced as described previously (*19, 20, 62–64*). A 3.0 mm-wide burr hole was made in the skull bone of anesthetized mice at a point midway between the lambda and bregma sutures and laterally midway between the central suture and temporalis muscle. Cortical ablation (For sensory motor cortex, 1.5 mm each from bregma to caudal and rostral, 3 mm to the right, and 1.0 mm in depth. For visual cortex, 1.5-4.5 mm to rostral, 3mm to the right from bregma) was performed by aspiration with a sterilized pipette. Sham-operated mice also received a burr hole without cortical ablation. Compared to a controlled cortical impact (CCI) model, this injury damages the brain tissue by a surgical ablation (contusion) but does not provide any blow or violent shaking (concussion).

#### Controlled Cortical Impact (CCI)

Moderate CCI was performed as previously described by (*29, 30, 65–67*). After mice were anesthetized, a 4.0-mm-wide burr hole was made in the skull bone at a point midway between the lambda and bregma sutures and laterally midway between the central suture and temporalis muscle. After removal of the scalp, the tip of the 3-mm impactor (Leica) piston was angled and kept perpendicular to the exposed cortical surface. The cortical impact was given with the following parameters: impact speed, 3.5 m/s; deformation depth, 1.0 mm; and duration, 400 ms. Sham mice underwent the same craniotomy without cortical impact.

#### Repeated closed CCI

Repeated closed CCI was performed as previously described (*68, 69*). Briefly, a midline incision was made to expose the skull. The 5-mm impactor piston tip was placed vertically 2 mm to the right of the bregma. The cortical impact was given with the following parameters: impact speed, 5.0 m/s; deformation depth, 1.0 mm; and duration, 200 ms. Sham mice underwent the same midline incision without cortical impact. Mice received one injury per day for three consecutive days.

### Cannula implantation

Cannula implantation and microinjection were done as previously described (*70–74*). 26-gauge guide cannula (Plastics One) was implanted ipsilaterally over the right side of the thalamus (−1.80 mm AP, +1.50 mm ML, and −3.00 mm DV) or right hippocampus (−1.80 mm AP, +1.50 mm ML, and −1.50 mm DV). The guide cannula was secured in place with anchoring miniature screws and dental cement. A 32-gauge dummy cannula (Plastics One) was inserted into each cannula to prevent clogging.

### Local antibody and LPS injection

Antibodies and LPS (4 µg/µl, L2880, Sigma) were administered via implanted cannula following the indicated schedule. The following antibodies and respective isotype controls were injected: anti-CSF1R (CD115) (3.0 µg/µl; Bio X Cell, #BE0213) and rat IgG2a isotype control (3.0 µg/µl; Bio X Cell, #BE0089); anti-CD9 (1.0 µg/µl; BD Biosciences, #553758) and normal rat IgG control (1.0 µg/µl; BD Biosciences, #553926); anti-FcγRIII (0.5 µg/µl; R&D Systems, #MAB19601) and normal rat IgG control (0.5 µg/µl; R&D Systems, #MAB006). On the injection day, the dummy cannula was replaced with a 33-gauge internal cannula (Plastics One) extending 2 mm below the tip of the guide cannula (Plastics One). Unless otherwise stated, 0.5 µl of antibodies or LPS were injected at 100 nl/min using a microinfusion pump (Narishige) per each injection. The internal cannula was left in place for an additional 5 min to diffuse antibodies or reagents properly.

### DREADD experiments

For neuronal DREADD experiments, AAV injection was conducted as previously described (*75*). Briefly, 1 µl of AAV-hSyn-hM3D (Gq) -mCherry (3.23×10^12^ GC/ml; Addgene, #50474) was stereotactically injected into the thalamus (−1.80 mm AP, +0.50 mm ML, and −3.20 mm DV) of WT mice using a 10 µl Nanofil syringe (WPI) at a rate of 200 nl/min immediately after cortical injuries. CNO (10 mg/kg; Cayman, 16882) or vehicle was administered intraperitoneally (i.p.) 30 min before the test session. For microglial DREADD experiments, *Tmem119 ^CreERT2^hM3Dq-mCitrine^LSL/LSL^*mice were injected with 0.5 µl CNO (0.5 mg/mL; Cayman, #16882) or vehicle at a rate of 100 nl/min via an implanted cannula using a microinfusion pump (Narishige) for three consecutive days before the analysis.

### Immunohistochemistry

Mice were anesthetized at two hours after the novel object recognition (NOR) test and transcardially perfused with ice-cold phosphate-buffered saline (PBS) (pH 7.4) and 4% paraformaldehyde (PFA)/PBS. Brain tissue was post-fixed in 4% PFA overnight, cryoprotected by incubating in 15% and 30% sucrose/PBS, and frozen in the O.C.T. compound. Free-floating sections (30 µm in thickness) were prepared by a Leica cryostat, placed in blocking buffer (5% normal goat serum and 0.1% Triton X-100 in PBS) for 1 hour at room temperature, and then incubated with primary antibodies overnight at 4°C. After washing in PBS, the sections were further incubated with 1:500 dilutions of fluorophore-conjugated secondary antibodies (Thermo Fisher Scientific) for 1 hour at room temperature, followed by DAPI staining (10 µg/ml; Thermo Fisher Scientific, D1306) for 10 min. The sections were mounted on slide glasses with ProLong Diamond antifade mounting medium (Thermo Fisher Scientific, P36961) or Fluorescent mounting media (DAKO, S3023). Images were acquired using Zeiss LSM 800 Airyscan confocal microscopes and a ZEN software (Zeiss). The following primary antibodies were used: rabbit polyclonal anti-Iba1 (1:500; WAKO, #019–19741,), Alexa Fluor 488-conjugated rabbit monoclonal anti-Iba1 (1:500; abcam, #ab225260), mouse monoclonal anti-Iba1 (1:500; Sigma-Aldrich, #MABN92), Alexa Fluor 647-conjugated mouse monoclonal anti-NeuN (1:500; abcam, #ab190565), rabbit monoclonal anti-NeuN (1:500; abcam, #ab177487), rabbit monoclonal anti-c-Fos (1:500; Cell Signaling Technology, #2250), rabbit monoclonal anti-TSPO (1:100; abcam, #ab109497), rat monoclonal anti-CD68 (1:2,000; Novus Biologicals, #NBP2-33337), mouse monoclonal anti-TNF-α (1:100; abcam, #ab1793), rat monoclonal anti-RFP (1:500; Chromotek, #5F8), rabbit monoclonal anti-HA-tag (1:500; Cell Signaling Technology, #3724), armenian hamster monoclonal anti-CD31/PECAM1 (1:100; DSHB, #2H8-s), rabbit polyclonal CD9 (1:100; Proteintech, #20597-1-AP), CD9 (1:100; BD Biosciences, #553758), rabbit polyclonal anti-FcγRIII (1:100; abcam, #ab203883), rabbit polyclonal anti-p-Syk (1:200; Cell Signaling, #2710T), mouse monoclonal anti-PSD95 (1:500; Thermo Fisher Scientific, #MA1-046), goat polyclonal anti-LPL (1:100; R&D Systems, #AF7197), chicken monoclonal anti-Tmem119 (1:500; Synaptic Systems, #400009), sheep polyclonal anti-Trem2 (1:100; R & D, #AF1729), rabbit polyclonal anti-Claudin5 (1:100, Thermo Fisher Scientific, #34-1600), and rabbit polyclonal anti-Ifitm3 (1:100; Proteintech, #11714-1-AP).

### Image analysis

Image analysis was performed as previously described (*75–77*). Z-stack images were used for evaluation. Image processing and subsequent quantifications were carried out on ImageJ (National Institute of Health). The “Measure Object Area” module was used to measure the Iba1 coverage area. The number of Iba1^+^, NeuN^+^, and c-Fos^+^ cells per each visual field was quantified using “Analyze Particles” module. To determine the coverage area of CD68^+^, p-Syk^+^, PSD95^+^, and IgG^+^ signals in Iba1^+^ cells, the boundaries of CD68^+^, p-Syk^+^, PSD95^+^, and IgG^+^ areas were traced and then superimposed onto the Iba1+ images using ImageJ. Then, the sizes of double/triple-positive areas were measured and divided by the number of Iba1^+^ microglia in the field to obtain the average size of CD68^+^, p-Syk^+^, PSD95^+^, and IgG^+^ areas per Iba1^+^ microglia. Then the data from three to five brain sections per mouse were averaged to represent each mouse. To evaluate the distribution of various microglia across the thalamus, we acquired 30 images from the right thalamus 21 days after cortical injury. The fluorescent intensities of Tmem119, CD9, and Ifitm3 signals in Iba1+ microglia were measured in each area. If the Tmem119 intensity was the highest, the area was defined as MG#2/3-rich area (green). Similarly, the areas with the highest CD9 and Ifitm3 signals were defined as MG#4 (blue) and MG#6-rich area (red). The area with similar intensities for CD9 and Tmem119 was defined as MG#1-rich area (yellow).

### Single-cell RNA sequencing (scRNA-seq)

#### Sample preparation and sequencing

Thalamic and hippocampus (HPC) tissues from the injured hemispheres were micro-dissected and single cell suspensions were obtained following the manufacturer’s protocol and as described previously (*78*). Briefly, tissue was transferred into 2 ml of pre-chilled homogenization buffer [15 mM HEPES buffer, 1 mg/mL DNase1 (Roche), and Protector RNase inhibitor at 1:200 (Roche)]. Tissues were carefully homogenized 10-12 times with Dounce homogenizers, filtered with 70 μm cell strainers, and centrifuged at 2,500 rpm for 2 min to pellet cells. Then, debris was removed using Debris Remove Solution (Miltenyi). Equal numbers of cells from 4 individual mice were pooled together for the thalamus and the HPC, respectively. Samples were Fc-blocked with anti-mouse CD16/32 antibody (1:100, Biolegend, clone: 93), and stained with 7-AAD (ThermoFisher), FITC-conjugated anti-mouse CD11b (1:100; Biolegend, clone: M1/70), and APC-conjugated anti-mouse CD45 (1:500; Biolegend, clone: 30-F11). Then, live CD45^+^ cells were sorted using the FACS Aria II at the UAB Comprehensive Flow Cytometry Core (CFCC). scRNA-seq libraries were constructed on the 10x Genomics platform using the Single Cell 3’ protocol (v.3.1, 10x Genomics). The libraries were sequenced using NovaSeq6000 at the UAB Genomics Core facility.

#### Data analysis

Raw scRNA-seq data were processed using the Cell Ranger software (v.6.0.2 10x Genomics) for sample demultiplexing, barcode processing, and single-cell counting. Reads were aligned to the GRCm38 (mm10) mus musculus reference genome. The outputs from the Cell Ranger software analysis were then imported into the Seurat package (v.4.0) in R (v.4.1.1), and the subsequent analysis was conducted with the Seurat. For quality control, cells containing more than 2% of mitochondrial counts and extreme unique feature counts (over 7,500 or less than 200) were removed. Data were normalized with a scaling factor of 10,000. The data from multiple libraries were then combined using FindIntegrationAnchors() and IntegrateData() functions. Principal component analysis (PCA) was performed using the combined data. A scree plot was generated using ElbowPlot() function to select the principal components (PCs) by localizing the last PC before the explained variance reaches a plateau. Selected PCs were used to calculate nearest-neighbor distances and perform Louvain clustering with FindNeighbors() and FindClusters() functions. Then, uniform manifold approximation and projection (UMAP) was used to visualize clusters. Cell types were initially annotated using SingleR (*79*), and then refined based on the expression of known marker genes in the genes significantly enriched for each cluster that was obtained using FindAllMarkers() function. Wilcoxon rank-sum test was used to identify genes differentially expressed between injury and sham groups per cluster. Among the 19 identified clusters, six clusters were annotated as microglia and used for further analysis. Gene set enrichment analysis (GSEA) was performed for the microglia cluster data using gsea() function of Clusterprofiler (v. 4.8.3) (*80, 81*). To visualize the GSEA result, Dotplot() and ggplot() functions were employed. Gene sets for homeostatic microglia (HM), disease-associated microglia (DAM), and interferon-responsive microglia (IRM) were created according to previous studies (*82–85*). To compare cluster compositions between the thalamus and HPC, cell number and normalized ratio in each cluster were counted. The result was visualized using ggplot() function. Individual gene expression patterns were visualized by FeaturePlot() function. Pseudo-time analysis was performed using Monocle 3 (*86–88*) on the six microglial clusters. These clusters were first randomly downsampled to 1000 cells per cluster by subset() function, and microglia cluster 3 was set as the pseudo-time root cluster based on the expression profile matching that of known homeostatic microglia. Pseudo-time was then calculated and plotted for visualization. A list of genes identified as important in calculating pseudo-time was also generated using Monocle 3. Individual gene expression patterns across cells were visualized by FeaturePlot() function. Findmarkers() function was used to identify the differentially expressed genes in MG#4 cluster between the thalamus and the HPC, and MG#4 clusters between the anti-CD9 Ab group thalamus and control Ab group thalamus. The results were visualized as a volcano plot using ggplot() function. The single-cell heatmap was generated using the DoHeatmap() function, while the pseudo-bulk heatmap was created by first calculating the average gene expression using the AggregateExpression() function, and then plotting the data using the ggplot() function. To minimize batch effects in the data obtained from experiments using antibody and in the comparison experiments between thalamus and hippocampus, the RunHarmony() function was used. The UMAP plot generated using the dataset created with the RunHarmony() function was randomly downsampled to 3,000 cells per condition by subset() function.

### Behavioral assays

The novel object recognition (NOR) test evaluates the tendency of rodents to discriminate between new and familiar objects and is used to assess recognition memory in mice (*89*). The test mice were individually habituated for 10 min in an open field box (45 x 45 centimeters; Harvard Apparatus) for two consecutive days. The next day, the test mice were exposed to the familiar open field with two identical objects placed at an equal distance in a symmetrical position for 10 min. Twenty-four hours later, the test mice were allowed to explore the same open field with the same object (familiar object) and a new object with a different shape (novel object). The exploratory behaviors of the test mice for 10 min were recorded and analyzed using EthoVision XT17 (Noldus). Exploration of the object was counted when the animal’s nose entered a 2 cm radius around the object. The discrimination index was calculated as ([time spent at the novel object] – [time spent at the familiar object])/([time spent at the novel object] + [time spent at the familiar object]).

### Statistical analysis

Data were analyzed with Student’s t test and one-way ANOVA using Microsoft Excel and GraphPad Prism 8 (GraphPad Software) unless stated otherwise. Post hoc analyses for one-way ANOVA were performed using Dunnett’s test method. Significant differences were considered at p < 0.05.

## List of Supplementary Materials

Table S1. List of genes whose expression was significantly upregulated in MG#1-6.

Table S2. Genes used for Cluster Profiler analysis.

Table S3. Outputs of pseudotime analysis with Monocle 3.

Table S4. List of differentially expressed genes between the thalamus and HPC MG#4.

Table S5. List of differentially expressed genes in MG#1 between control Ab- and anti-CD9 Ab-treated mice.

Table S6. List of differentially expressed genes in MG#4 between control Ab- and anti-CD9 Ab-treated mice.

Table S7. Statistical analyses used in this study.

## Acknowledgments

We thank Dania Mallah and Vidhula Prasanna for their technical help; the members of Kano and Niwa labs for their critical discussion on this manuscript; Shanrun Liu (UAB Flow Cytometry and Single-Cell Core Facility) for single-cell RNA-seq library preparation; Michael Crowley (UAB Heflin Center Genomics Core Lab) for Illumina sequencing. We acknowledge the following funding support: National Institutes of Health (NIH) (R01NS133242) and a UAB 1R01 award to S.K.; a Uehara Memorial Foundation Overseas Postdoctoral Fellowship to K.M.; a Japan Society for the Promotion of Science (JSPS) Postdoctoral Fellowship to K.T.; and NIH (T32GM008361, T32MH129274, and F30MH139313) to J.B. The UAB Flow Cytometry and Single Cell Services Core is supported by NIH grants to the O’Neal Comprehensive Cancer Center (P30CA013148) and the Center for AIDS Research (P30AI027767). S.K. is an Allen Distinguished Investigator, a Allen Frontiers Group advised program of the Paul G. Allen Family Foundation (now Allen Family Philanthropies).

## Supplementary Materials

**Fig. S1.**
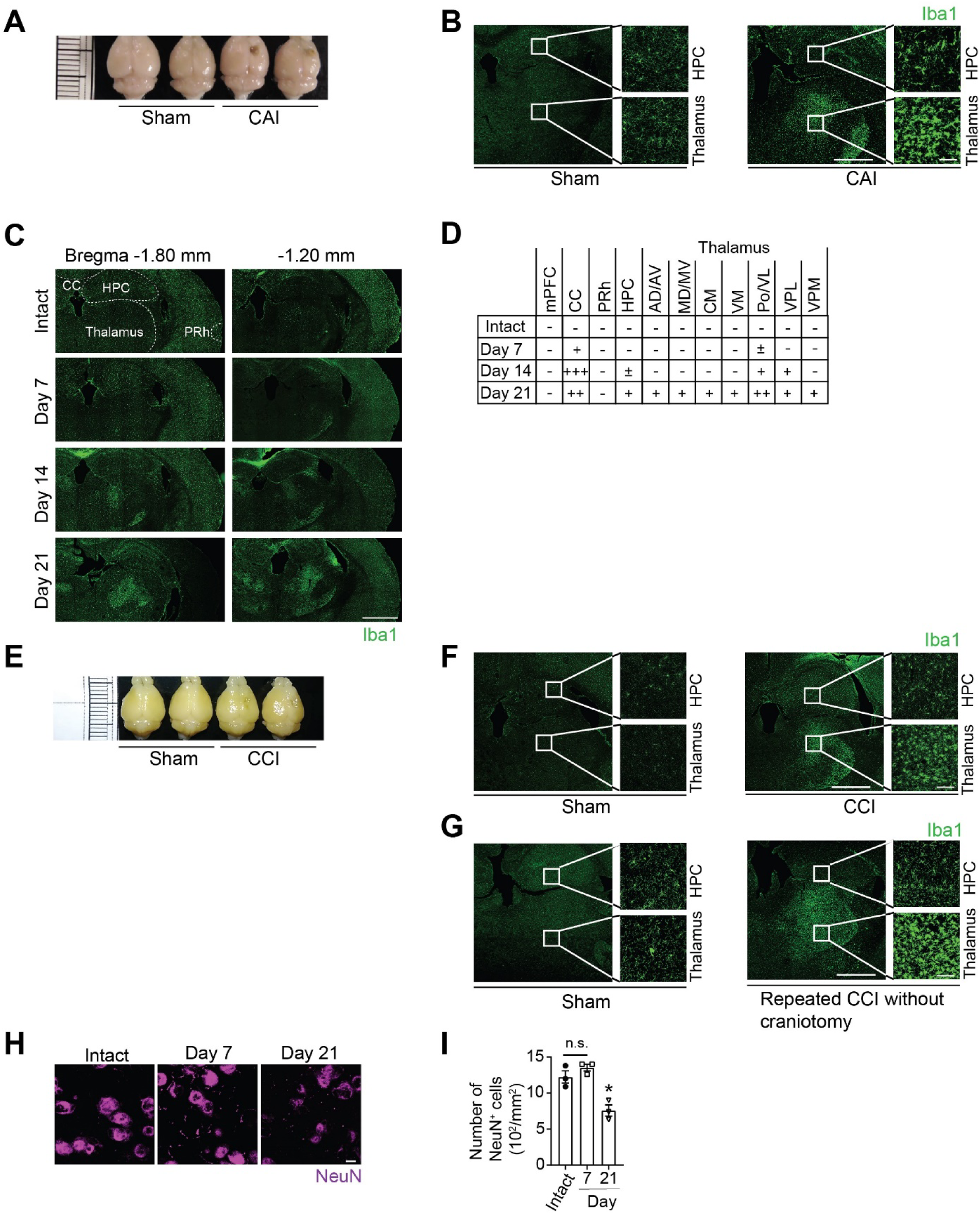
Cortical brain injuries induce delayed microglia activation. (A) Representative images of the gross brain morphology 21 days after the unilateral cortical ablation injury (CAI) and sham surgery. (B) Representative images of Iba1^+^ microglia 21 days after CAI and sham surgery. Scale bars: 1 mm (in low-magnification image) and 100 µm (in high-magnification image). HPC, hippocampus. (C) Representative images of Iba1^+^ microglia in the intact brain and the brains on day 7, 14, and 21 post-CAI. Scale bar: 500 µm. CC, corpus callosum; PRh, perirhinal cortex. (D) Summary of delayed microglial reactive changes across various brain regions. The severity of microglia activation is indicated as follows: -, no activation; ±, faint activation; +, mild activation; ++, moderate activation; and +++, massive activation. mPFC, medial prefrontal cortex; CC, corpus callosum; PRh, perirhinal cortex; HPC, hippocampus; AD, anterodorsal nucleus; AV, anteroventral nucleus; MD, mediodorsal nucleus; MV, medioventral nucleus; CM, central medial nucleus; VM, ventral medial nucleus; VL, ventral lateral nucleus; Po, posterior nucleus; VPL, ventral posterolateral nucleus; VPM, ventral posteromedial nucleus. (E) Representative images of the gross brain morphology 21 days after controlled cortical impact (CCI) and sham surgery. (F) Representative images of Iba1^+^ microglia 21 days after open CCI and sham surgery. Scale bars: 1 mm (in low-magnification image) and 100 µm (in high-magnification image). (G) Representative images of Iba1^+^ microglia 21 days after repeated closed CCI and sham surgery. Scale bars: 1 mm (in low-magnification image) and 100 µm (in high-magnification image). (H) Representative images of NeuN^+^ neurons in the intact thalamus and 7 and 21 days after injuries. Scale bar, 10 µm. (I) Quantification data of the number of NeuN^+^ neurons in the thalamus (n = 3 mice per time point). In bar graphs, data are shown as the mean□±□s.e.m, and each dot/rectangle represents an individual animal. \*\*\**p* < 0.001; Student’s *t*-test.

**Fig. S2.**
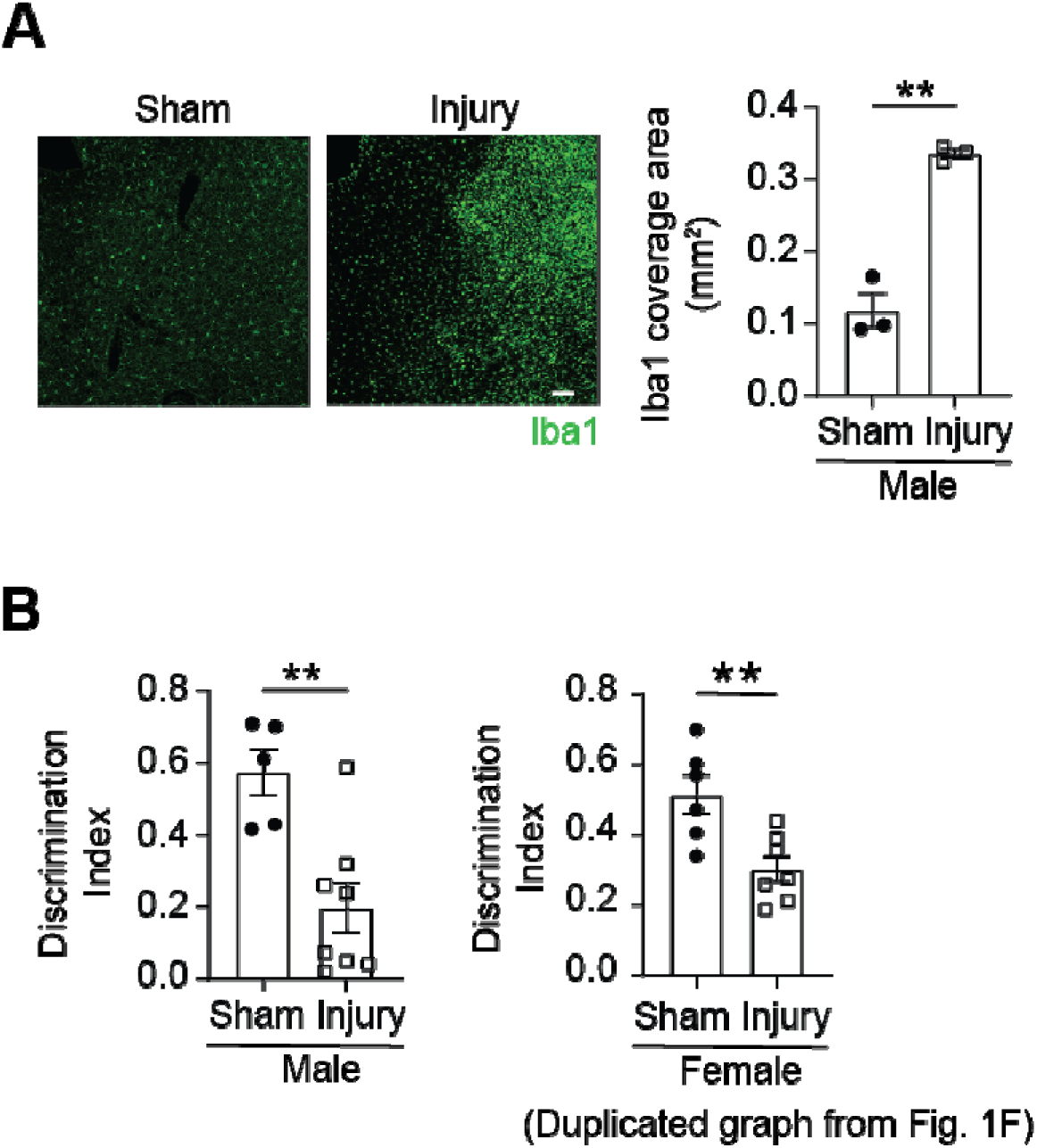
Cortical brain injuries in male mice cause thalamic microglial reactive changes and the NOR deficits similar to those in female mice 21 days after. (A) Representative images of Iba1^+^ microglia in the thalamus 21 days after injuries and sham surgery. Scale bar: 100 µm. Quantification of Iba1^+^ area is shown (n = 3 mice per group). (B) Discrimination index in the NOR test for sham (n = 5 mice) and injury (n = 8 mice) group. Female data on the right were duplicated from Fig. 1F for reference. In bar graphs, data are shown as the mean□±□s.e.m, and each dot/rectangle represents an individual animal. \*\**p* < 0.01; Student’s *t*-test.

**Fig. S3.**
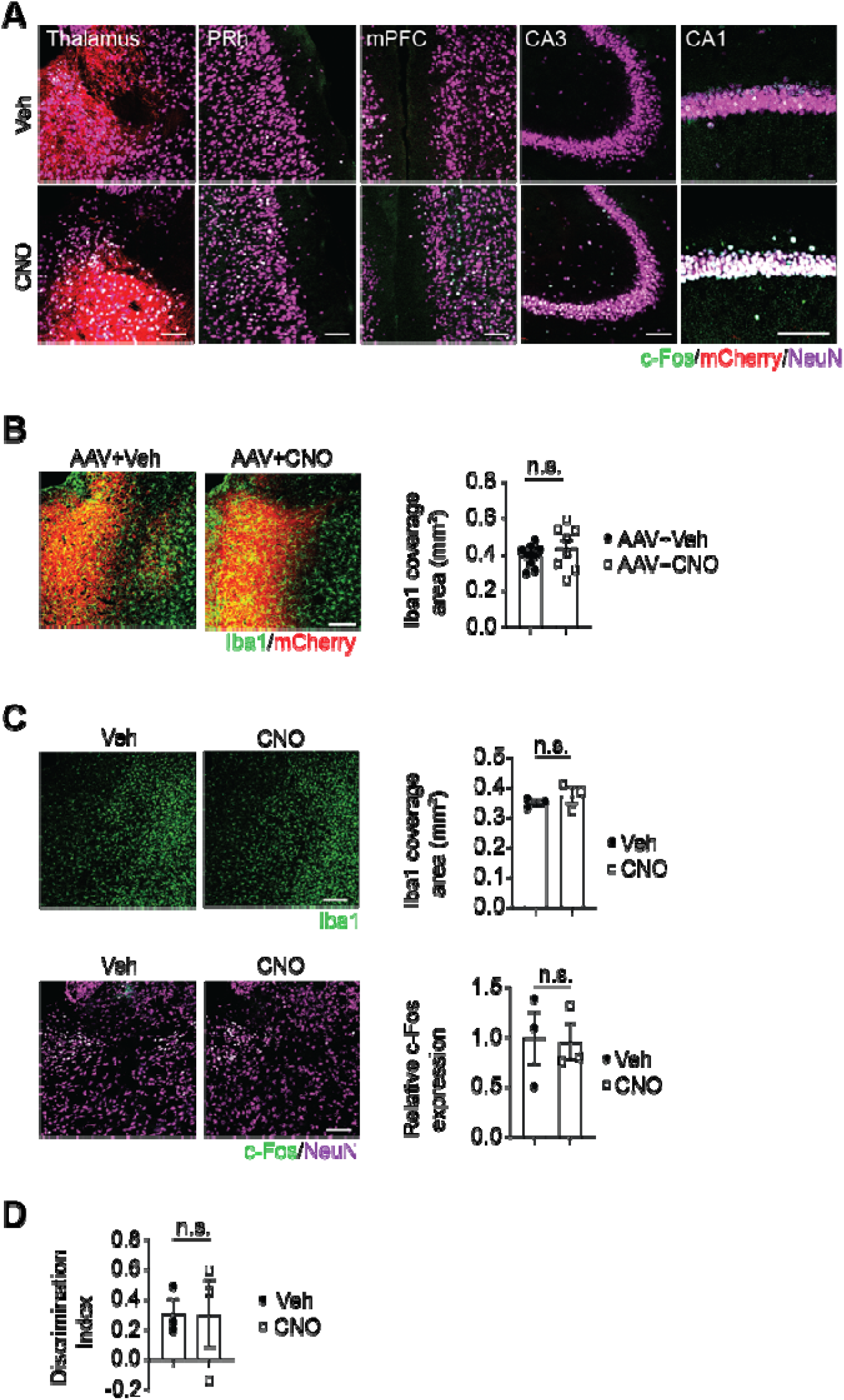
Chemogenetic activation of thalamic neurons ameliorate the NOR deficits after cortical injuries. (A) Representative images of neuronal c-Fos expression in the thalamus, PRh, mPFC, and HPC (CA3 and CA1) upon CNO or vehicle administration. (B) Representative images of Iba1^+^ microglia in AAV-injected areas (indicated by mCherry expression) in the thalamus of mice with CNO or vehicle administration. Quantification of Iba1 coverage area is shown on the right (n = 10) mice for AAV with vehicle; n = 8 mice for AAV with CNO). (C) Effects of CNO injection on thalamic microglia and neurons in injured mice without AAV injection. Upper panel: representative images of Iba1^+^ microglia in the thalamus after CNO or vehicle administration. Quantification of Iba1 coverage area is shown on the right (n = 3 mice per group). Lower panel: representative images of neuronal c-Fos expression in the thalamus after CNO or vehicle administration. Quantification is shown on the right (n = 3 mice per group). (D) Discrimination index in the NOR test for vehicle and CNO-treated injured mice (n = 3 mice per group). Scale bars: 100 µm (A, B, and C). In bar graphs, data are shown as the mean□±□s.e.m, and each dot/rectangle represents an individual animal. n.s., not significant; Student’s *t*-test.

**Fig. S4.**
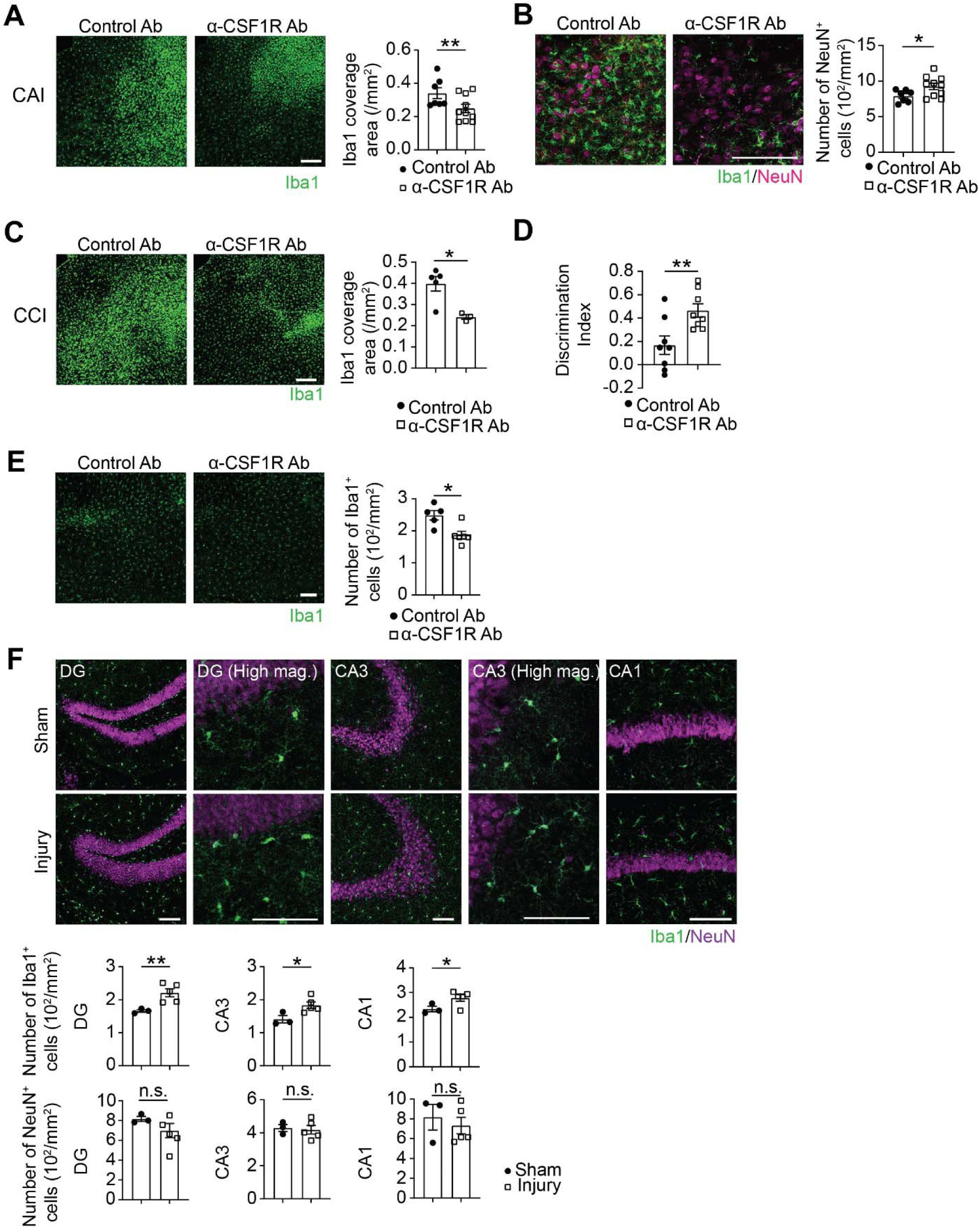
Microglial reactive changes and their contributions to NOR deficits differ between the thalamus and HPC in the injured mice. (A) Representative images of Iba1^+^ microglia in the thalamus of mice receiving intrathalamic injections of control Ab or anti-CSF1R Ab after cortical injuries (CAI). Quantification data of the Iba1 coverage area are shown on the right (n = 7 mice for control Ab; n = 10 mice for anti-CSF1R Ab). (B) Representative images of NeuN^+^ neurons in the thalamus of mice receiving intrathalamic injections of control Ab or anti-CSF1R Ab after cortical injuries. Quantification data of the number of NeuN^+^ cells are shown on the right (n = 7 mice for control Ab; n = 10 mice for anti-CSF1R Ab). (C) Representative images of Iba1^+^ microglia in the thalamus of mice receiving intrathalamic injections of control Ab or anti-CSF1R Ab after cortical injuries (CCI). Quantification data of the Iba1 coverage area are shown on the right (n = 5 mice for control Ab; n = 3 mice for anti-CSF1R Ab). (D) Discrimination index in the NOR test for mice with intrathalamic injections of control Ab and anti-CSF1R Ab (n =8 mice per group) after cortical injuries (CCI). (E) Representative images of Iba1^+^ microglia in the HPC of mice receiving intrahippocampal injections of control Ab or anti-CSF1R Ab after cortical injuries (CAI). Quantification data of the number of Iba1^+^ microglia are shown on the right (n = 5 mice for control Ab; n = 6 mice for anti-CSF1R Ab). (F) Representative images of Iba1^+^ microglia and NeuN^+^ neurons in the HPC 21 days after cortical injuries (CAI) and sham surgeries. Quantification data for each region are shown on the right (n = 3 mice for sham; n = 5 mice for injury). Scale bars: 100 µm. In bar graphs, data are shown as the mean□±□s.e.m, and each dot/rectangle represents an individual animal. n.s., not significant; \**p* < 0.05; \*\**p* < 0.01; Student’s *t*-test.

**Fig. S5.**
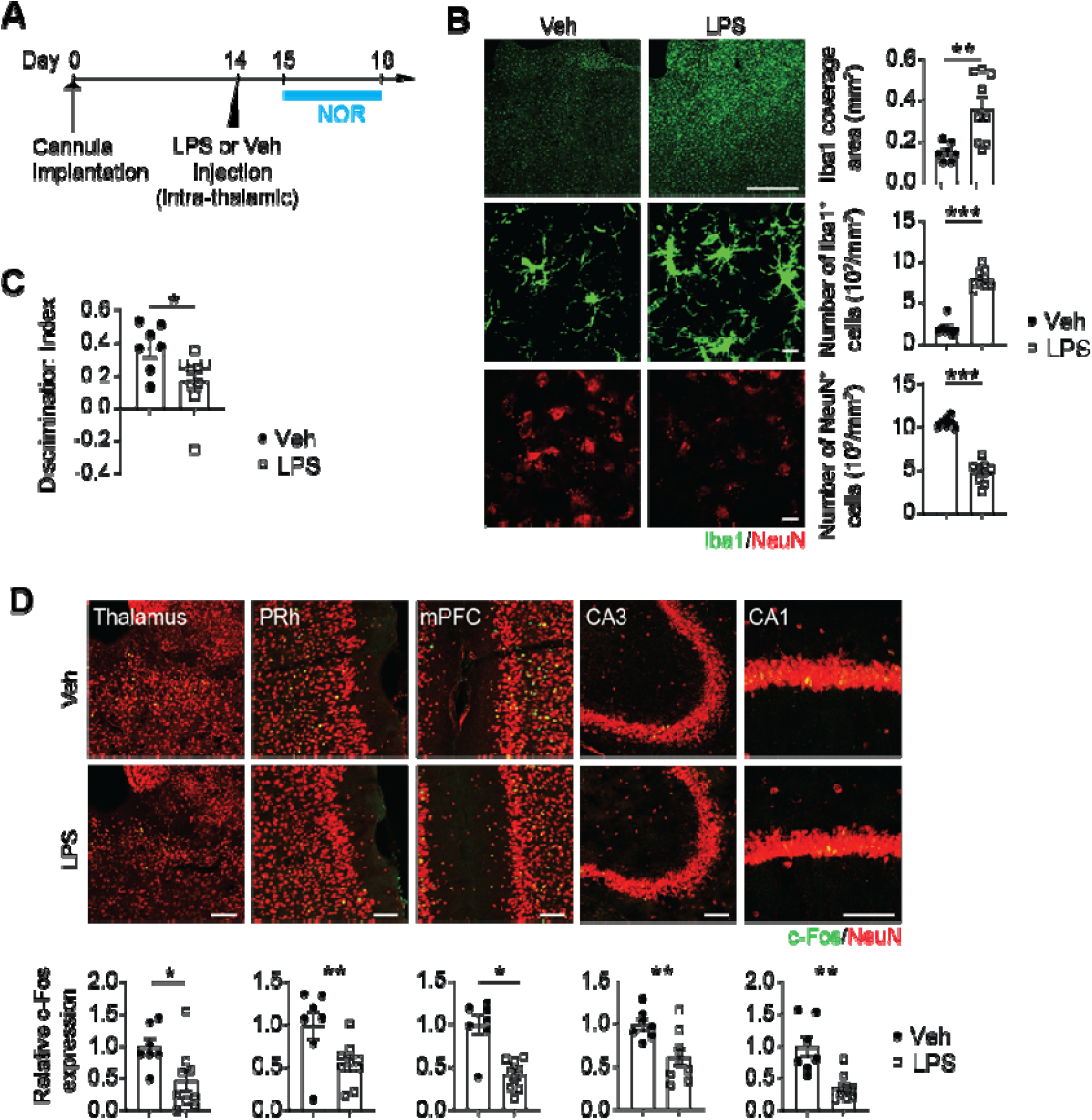
Reactive microglial changes induced by intra-thalamic LPS injection cause cognitive impairment in non-injured mice. (A) Experimental timeline for cannula implantation, intra-thalamic LPS injection, and the NOR test. (B) Representative images of Iba1^+^ microglia and NeuN^+^ neurons in the thalamus 4 days after vehicle or LPS injection. Quantification data of the Iba1 coverage area, the number of Iba1^+^ microglia, and the number of NeuN^+^ neurons are shown on the right (n = 7 and 9 mice for the vehicle- and LPS-injected group, respectively). Scale bars: 500 µm (top panel) and 10 µm (middle and bottom panels). (C) Discrimination index in the NOR test (n = 7 and 9 mice for the vehicle- and LPS-injected group, respectively). (D) Neuronal c-Fos expression in the thalamus, PRh, mPFC, and HPC (CA3 and CA1) 4 days after vehicle or LPS injection. Scale bars: 100 µm. In bar graphs, data are shown as the mean□±□s.e.m, and each dot/rectangle represents an individual animal. \**p* < 0.05; \*\**p* < 0.01; \*\*\**p* < 0.001; Student’s *t*-test.

**Fig. S6.**
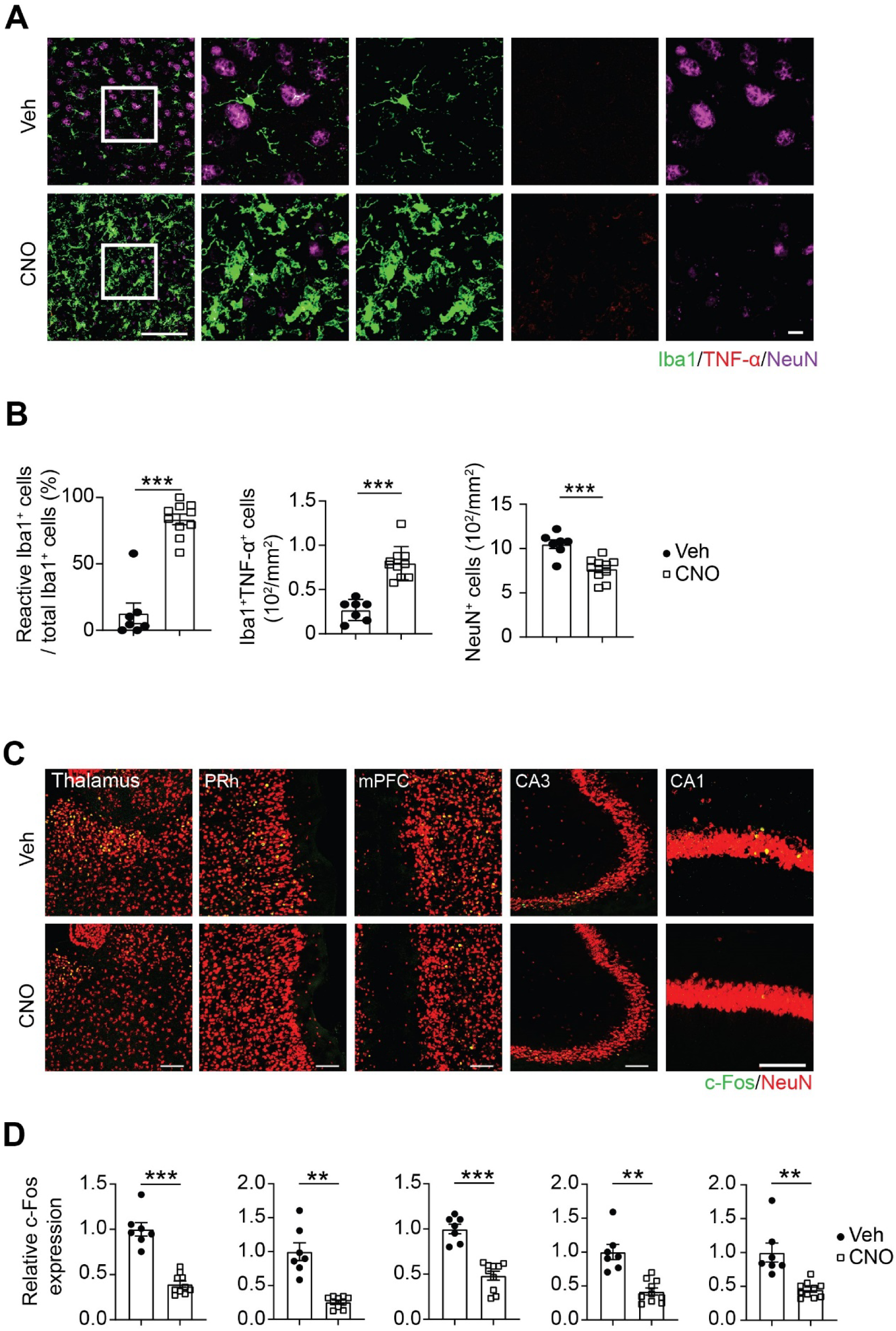
Chemogenetic induction of reactive changes in thalamic microglia causes cognitive impairment in non-injured mice. (A) Representative images of TNF-α producing Iba1^+^ microglia and NeuN^+^ neurons in the thalamus of the non-injured mice with vehicle or CNO injection. Scale bars: 100 µm (left panel) and 10 µm (right panel). (B) Quantification data of % microglia with reactive changes, TNF-α producing Iba1^+^ microglia, and NeuN^+^ cells (n = 7 mice for the vehicle group; n = 10 mice for the CNO group). (C) Representative images of neuronal c-Fos expression in the thalamus, PRh, mPFC, and HPC (CA3 and CA1) after vehicle or CNO injection. Scale bars: 100 µm (D) Quantification data of neuronal c-Fos expression changes in the CNO-treated group relative to the vehicle-treated group (n = 7 mice for the vehicle group; n = 10 mice for the CNO group). In bar graphs, data are shown as the mean□±□s.e.m, and each dot/rectangle represents an individual animal. \**p* < 0.05; \*\**p* < 0.01; \*\*\**p* < 0.001; Student’s *t*-test.

**Fig. S7.**
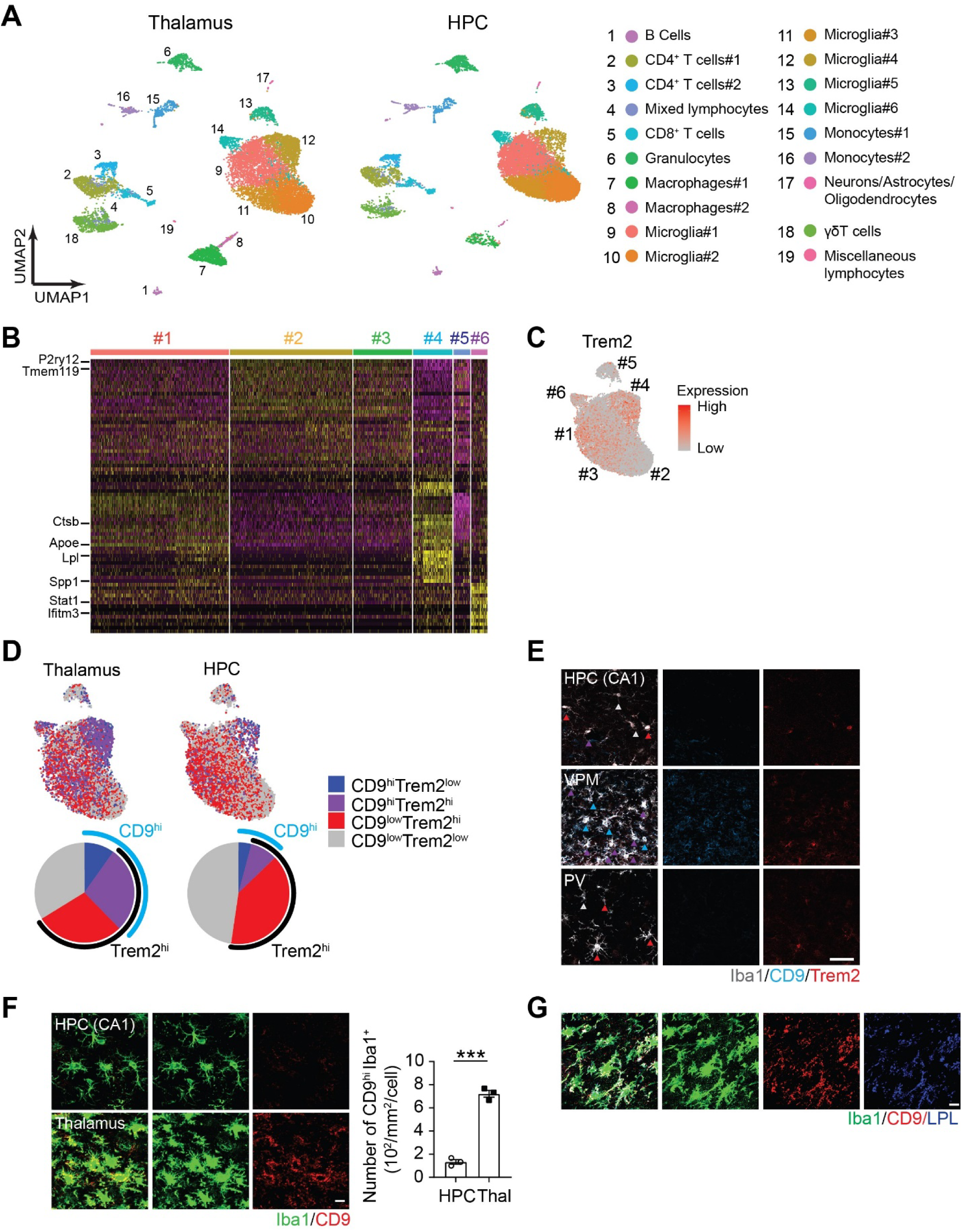
Microglia with distinct transcriptional states exist in the thalamus and HPC. (A) UMAP plot showing 19 clusters with 6 microglia subclusters in the hippocampus and the thalamus after cortical injury. (B) Heatmap showing unbiased top markers defining each microglia subtype after cortical brain injuries. (C) Feature plot showing Trem2 expression in all microglia clusters of the thalamus and HPC after cortical brain injuries. (D) Feature plots showing CD9 (blue) expression and Trem2 (red) expression in all microglia clusters of the thalamus and HPC after cortical brain injuries. Pie charts showing the proportion of CD9^hi^Trem2^lo^, CD9^hi^Trem2^hi^, CD9^lo^Trem2^hi^, and CD9^lo^Trem2^lo^ microglia clusters of the thalamus and HPC after cortical brain injuries. (E) Spatial distribution of CD9^hi^ microglia and Trem2^hi^ microglia in the thalamus after cortical brain injuries. Representative images of CD9^hi^ Trem2^hi^ microglia (purple arrowheads), CD9^hi^ Trem2^low^ microglia (blue arrowheads), CD9^low^ Trem2^hi^ microglia (red arrow heads), and CD9^low^ Trem2^low^ microglia (gray arrowheads) in the HPC, ventral posteromedial nucleus (VPM), and paraventricular nucleus (PV). Scale bar: 50 µm. (F) Representative images of CD9^hi^ Iba1^+^ cells in the HPC and thalamus after cortical brain injuries. Scale bar: 10 µm. Quantification of the number of CD9^hi^ Iba1^+^ microglia in the HPC (n = 3) and the thalamus (n = 3) is shown in bar graphs. (G) Representative images of Iba1^+^ microglia of the thalamus after cortical brain injuries that express CD9 and LPL. Scale bar: 10 µm. In bar graphs, data are shown as the mean□±□s.e.m, and each dot/rectangle represents an individual animal. \*\*\**p* < 0.005; Student’s *t*-test.

**Fig S8.**
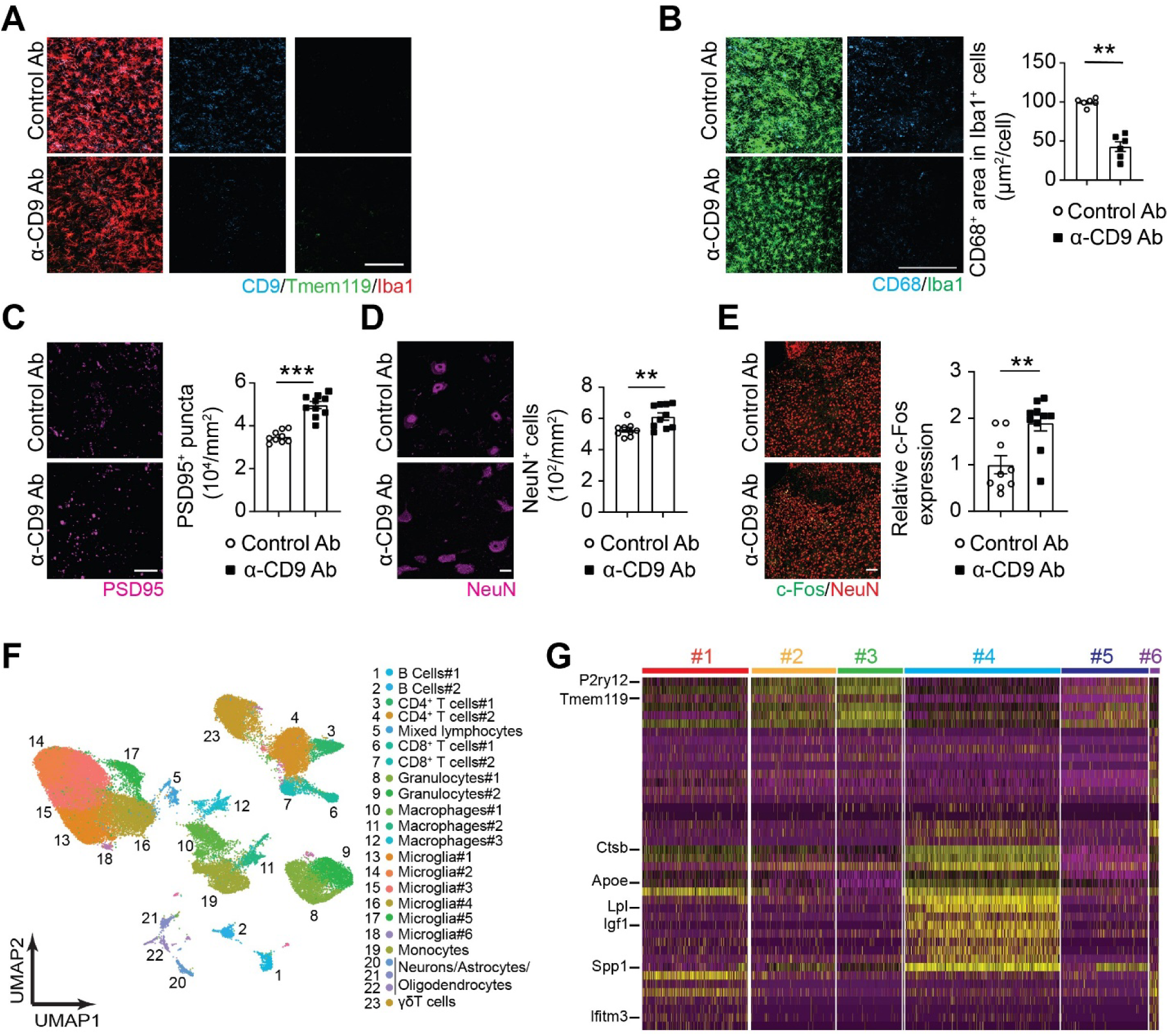
CD9 blockade inhibits MG#4 induction in the thalamus in the cortically injured mice. (A) Representative images of CD9^hi^ Iba1^+^ cells and Tmem119^hi^ Iba1^+^ cells in the thalamus in control Ab- and anti-CD9 Ab-injected mice after cortical brain injuries. Scale bar: 100 µm. (B) Representative images of CD68^+^ lysosomes in Iba1^+^ microglia in the thalamus in control Ab and anti-CD9 Ab-injected group after cortical brain injuries. Quantification data of average CD68^+^ area per microglia are shown in bar graphs for the control Ab- (n= 9 mice) and the anti-CD9 Ab-injected group (n = 10 mice). Scale bar: 100 µm. (C) Representative images of PSD95^+^ synaptic puncta of the thalamus in control Ab- and anti-CD9 Ab-injected mice after cortical brain injuries. Quantification data are shown in bar graphs for the control Ab- (n = 9 mice) and the anti-CD9 Ab-injected group (n = 10 mice). Scale bar: 10 µm. (D) Representative images of NeuN^+^ cells of the thalamus in control Ab- and anti-CD9 Ab-injected mice after cortical brain injuries. Quantification data are shown in bar graphs for the control Ab- (n = 9 mice) and the anti-CD9 Ab-injected group (n = 10 mice). Scale bar: 10 µm. (E) Representative images of neuronal c-Fos expression in the thalamus in control Ab- and anti-CD9 Ab-injected mice after cortical brain injuries. Quantification data of neuronal c-Fos expression in the anti-CD9 Ab group (n = 10 mice) relative to the control Ab group (n = 9 mice) are shown in bar graphs. Scale bar: 100 µm. (F) UMAP plot showing 23 clusters with 6 microglia subclusters in the thalamus of cortically injured mice injected with control and anti-CD9 Ab. (G) Heatmap showing unbiased top markers defining each microglia subtype in the thalamus after control Ab- and anti-CD9 Ab-injection. In bar graphs, data are shown as the mean□±□s.e.m, and each dot represents an individual animal. n.s., not significant; \*\**p* < 0.01, \*\*\**p* < 0.005; Student’s *t*-test.

**Fig. S9.**
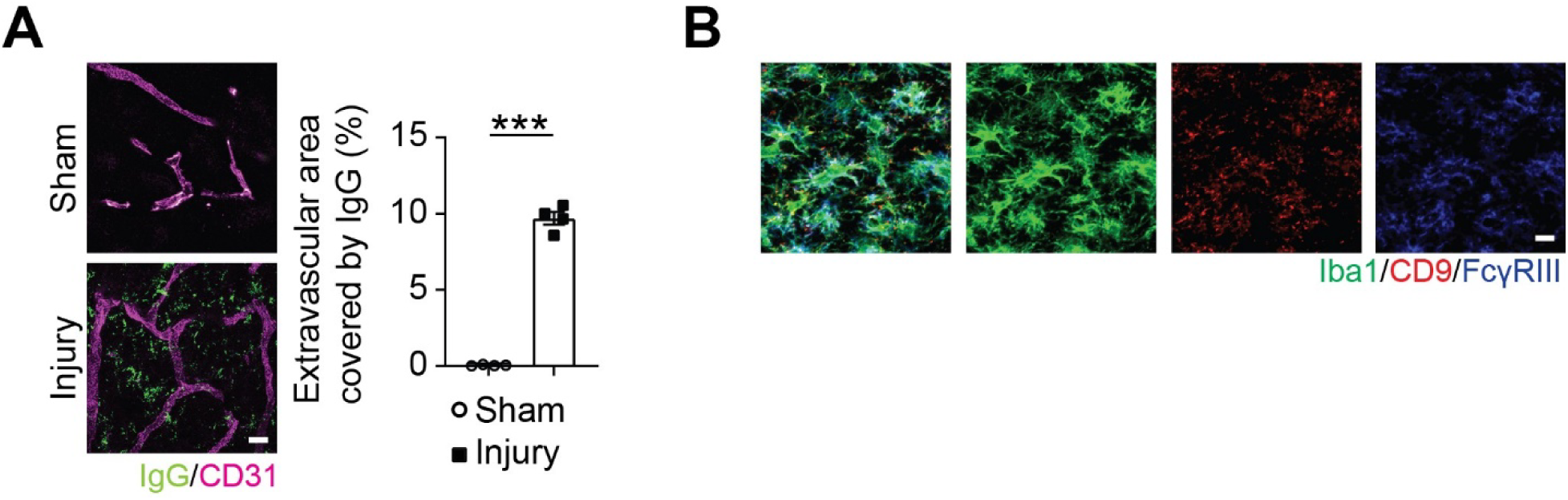
Extravasated IgG and FcγRIII expression in CD9hi microglia are observed in the thalamus in the cortically injured mice. (A) Representative images of IgG and CD31^+^ blood vessels in the thalamus after cortical brain injuries and sham surgery. Percentages of extravascular IgG coverage area per total IgG coverage area are shown in bar graphs for the injury (n = 4 mice) and the sham (n = 4 mice) group. Scale bar: 10 µm. (B) Representative images of CD9 and FcγRIII expression in the same Iba1^+^ microglia. Scale bar: 10 µm. In bar graphs, data are shown as the mean□±□s.e.m, and each dot represents an individual animal. \*\*\**p* < 0.001; Student’s *t*-test

